# Global stress response in *Pseudomonas aeruginosa* upon malonate utilization

**DOI:** 10.1101/2024.03.26.586813

**Authors:** Karishma Bisht, Moamen M. Elmassry, Hafij Al Mahmud, Shubhra Bhattacharjee, Amrika Deonarine, Caroline Black, Michael J. San Francisco, Abdul N. Hamood, Catherine A. Wakeman

## Abstract

Versatility in carbon source utilization assists *Pseudomonas aeruginosa* in its adaptation to various niches. Recently, we characterized the role of malonate, an understudied carbon source, in quorum sensing regulation, antibiotic resistance, and virulence factor production in *P. aeruginosa*. These results indicate that global responses to malonate metabolism remain to be uncovered. We leveraged a publicly available metabolomic dataset on human airway and found malonate to be as abundant as glycerol, a common airway metabolite and carbon source for *P. aeruginosa*. Here, we explored and compared adaptations of *P. aeruginosa* UCBPP-PA14 (PA14) in response to malonate or glycerol as a sole carbon source using transcriptomics and phenotypic assays. Malonate utilization activated glyoxylate and methylcitrate cycles and induced several stress responses, including oxidative, anaerobic, and metal stress responses associated with increases in intracellular aluminum and strontium. Some induced genes were required for optimal growth of *P. aeruginosa* in malonate. To assess the conservation of malonate-associated responses among *P. aeruginosa* strains, we compared our findings in strain PA14 with other lab strains and cystic fibrosis isolates of *P. aeruginosa*. Most strains grew on malonate as a sole carbon source as efficiently as or better than glycerol. While not all responses to malonate were conserved among strains, formation of biomineralized biofilm-like aggregates, increased tolerance to kanamycin, and increased susceptibility to norfloxacin were the most frequently observed phenotypes. Our findings reveal global remodeling of *P. aeruginosa* gene expression during its growth on malonate as a sole carbon source that is accompanied by several important phenotypic changes. These findings add to accumulating literature highlighting the role of different carbon sources in the physiology of *P. aeruginosa* and its niche adaptation.

**Importance:** *Pseudomonas aeruginosa* is a notorious pathogen that causes local and systemic infections in immunocompromised individuals. Different carbon sources can uniquely modulate metabolic and virulence pathways in *P. aeruginosa*, highlighting the importance of the environment that the pathogen occupies. In this work, we used a combination of transcriptomic analysis and phenotypic assays to determine how malonate utilization impacts *P. aeruginosa,* as recent evidence indicates this carbon source may be relevant to certain niches associated within the human host. We found that malonate utilization can induce global stress responses, alter metabolic circuits, and influence various phenotypes of *P. aeruginosa* that could influence host colonization. Investigating the metabolism of malonate provides insight into *P. aeruginosa* adaptations to specific niches where this substrate is abundant, and how it can be leveraged in the development of much-needed antimicrobial agents or identification of new therapeutic targets of this difficult-to-eradicate pathogen.

## Introduction

*Pseudomonas aeruginosa* is an opportunistic pathogen that infects various hosts including humans [1]. It can infect a wide range of body sites, e.g., lungs in cystic fibrosis patients, wounds and blood circulation in burn and trauma patients [2]. Its versatile metabolism enables *P. aeruginosa* to survive quite different environmental conditions by utilizing a multitude of carbon sources [3]. This metabolic versatility is controlled by complex regulatory networks [4]. The utilization of certain carbon sources is linked to antibiotic tolerance and altered virulence of *P. aeruginosa* [5–7].

Furthermore, the activation of metabolic networks by specific carbon sources involves a stress response depending on the environmental conditions [8, 9]. However, the link between carbon source utilization and stress response is not clear.

Several microorganisms can utilize malonate, a naturally occurring organic acid in the human body and environment, as a carbon source [6, 10–13]. Previous studies suggest the potential clinical importance of malonate. For example, malonate utilization genes are overexpressed in *P. aeruginosa* grown ex vivo in blood from trauma patients [6]. Moreover, malonate increases the tolerance of *P. aeruginosa* to aminoglycoside antibiotics and regulates its quorum sensing circuit and virulence factors [5, 6, 14].

These malonate-induced shifts in metabolism were also associated with the production of surface-free biofilm-like aggregates (also known as flocs) embedded in a biomineralized matrix [14]. Similar bacterial aggregates have been associated with biofilms and lung infections [15, 16]. However, it is not known how carbon source utilization is involved in this phenotype.

Malonate utilization in *P. aeruginosa* not only triggers the previously mentioned adaptations, but also enhances the production of catalase, a possible indication to oxidative stress [14]. Yet, it is not known how malonate utilization is linked to oxidative stress response. Therefore, we hypothesized that malonate utilization induces a global stress response in *P. aeruginosa*, thus altering its phenotype. Using transcriptomics, we found that malonate utilization induces metals, oxidative, and other stress responses in *P. aeruginosa*. This was associated with the intracellular accumulation of certain metal ions. Next, we identified the genetic requirements for *P. aeruginosa* growth using malonate as a sole carbon source. Furthermore, some aspects of the malonate utilization-associated phenotypes were consistent among the tested lab strains and clinical isolates. Our findings highlight the lifestyle modulation effect of carbon sources on *P. aeruginosa* and provide evidence of malonate-inducing stress response and its associated phenotypes in this pathogen.

## Results

### Malonate as an abundant metabolite in human upper and lower airway

The human airway is a complex environment with many metabolites that can serve as sources of energy for *P. aeruginosa*. Among these is glycerol, which is an abundant metabolite and a degradation product of phosphatidylcholine, a major lung surfactant. Other groups and we have previously shown that malonate can be used as a carbon source and can alter *P. aeruginosa* phenotypes. Yet, despite *P. aeruginosa* being a common airway pathogen it has not been established whether malonate is relevant to this specific niche and if it is an available metabolite. To this end, we utilized a publicly available data set [17], where the authors sampled 8 healthy volunteers using four sampling methods (32 samples in total) to comprehensively examine the metabolites of the upper and lower airways using Ultra-high-Performance Liquid Chromatography-Tandem Mass Spectrometry (UPLC-MS/MS).

We aimed here to take an unbiased approach. Therefore, from the 581 detected metabolites, we filtered metabolites that are present in all 32 samples and confirmed using chemical standards. Following this selection, 160 metabolites remained, which included well-established metabolites in the airways (**Supplementary** Figure 1A).

Malonate was among these metabolites and found at a level similar to glycerol in all samples (**Supplementary** Figure 1B). While many other potential carbon sources were also detected, which is beyond the scope of this work, we were motivated by this observation and our previous results on how malonate alters *P. aeruginosa* phenotypes, e.g., virulence factors production and antibiotic resistance. Therefore, we aimed next to decipher how malonate utilization alters *P. aeruginosa* at the transcriptional level.

### Malonate utilization activates the glyoxylate and methylcitrate cycles

We sought to understand how malonate utilization alters the transcriptional regulatory circuits of carbon metabolism in *P. aeruginosa.* To uncover this, we examined the transcriptomic changes in *P. aeruginosa* strain UCBPP-PA14 (PA14) when grown in M9, a minimal medium, containing malonate (MM9), as a sole carbon source, vs. glycerol as a control (GM9). Transcriptomic data analysis was performed using the Rockhopper 2 [18]. Overall, 2,478 genes were differentially expressed (Benjamini-Hochberg procedure-adjusted P-value ≤ 0.05). In comparison to GM9, MM9 induced the expression of 1,370 genes and downregulated the expression of 1,108 genes. We validated our transcriptomic results, obtained from RNA-seq analysis, by examining the expression of a subset of genes of interest using qRT-PCR (Supplementary Figure 2).

The gene expression data enabled us to establish which metabolic pathways were activated by the growth of PA14 on malonate as a carbon source. As expected, the growth of *P. aeruginosa* in minimal medium with only malonate supplemented as a sole carbon source upregulated the expression of the 10 genes involved in malonate utilization, i.e., regulation, uptake, and decarboxylation **(Supplementary** Figure 3). The average expression ratio of these malonate utilization-related genes was 135 times higher in MM9 than in GM9.

Next, we identified 534 differentially expressed genes mapped to KEGG metabolic pathways, of which 70 mapped to carbon metabolism. Specifically, over 50 genes were mapped to pathways involving tricarboxylic acid (TCA), glyoxylate, and methylcitrate cycles **(Figure 1).** Upon utilization of malonate, acetyl-CoA is produced, which then feeds carbon metabolic circuits. Unlike the inhibited TCA cycle genes, the genes of glyoxylate and methylcitrate cycles were activated upon malonate utilization. These findings provide insight into how malonate utilization regulates different carbon metabolic pathways.

**Figure 1.**
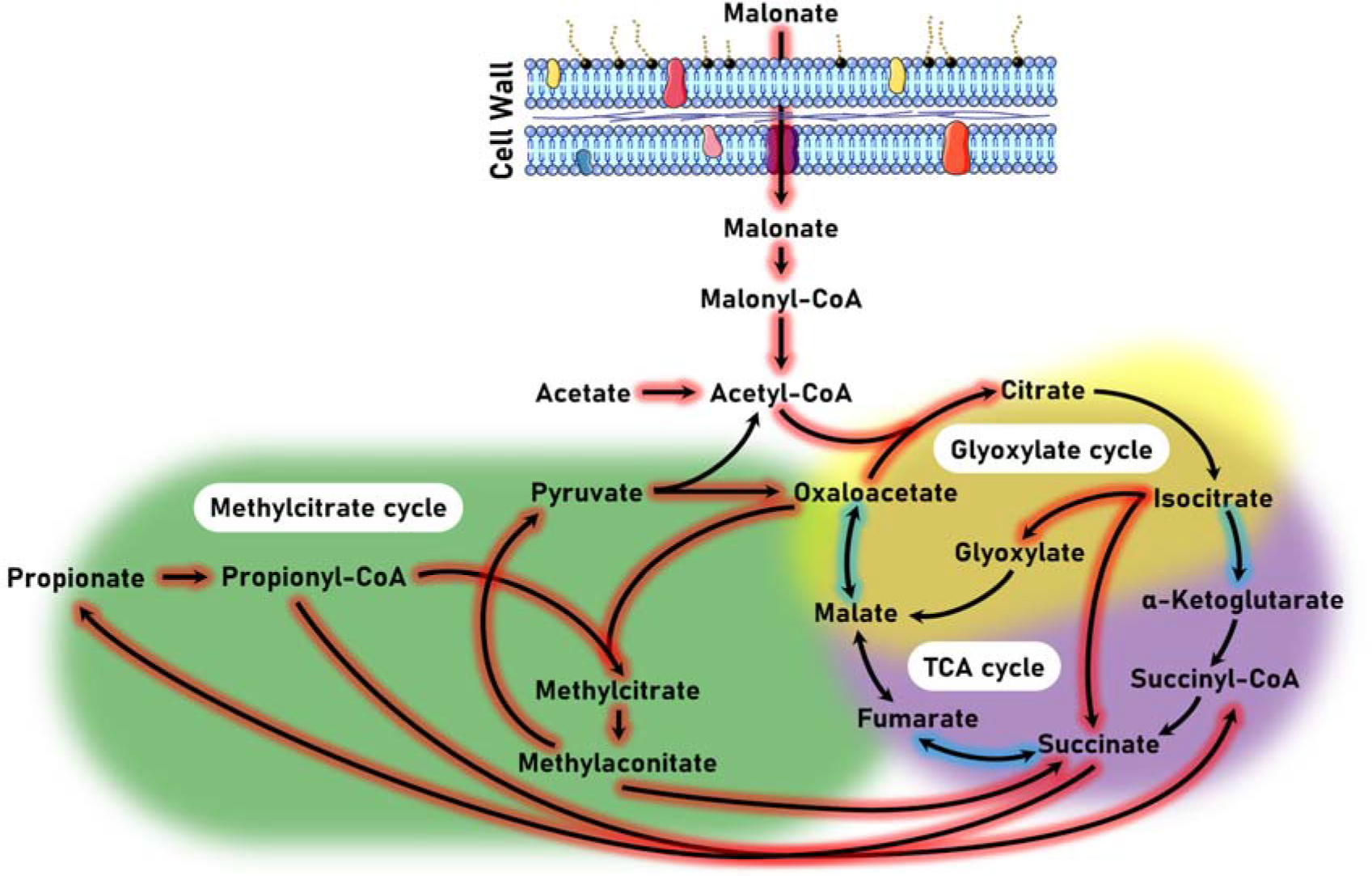
Transcriptomic analysis revealed the differential regulation of genes associated with different metabolic cycles. Central carbon metabolic pathways involved in malonate utilization in *P. aeruginosa*. Arrows represent the differentially expressed genes and their corresponding reactions. Red-highlighted arrows indicate upregulation and blue highlighting indicates downregulation. The three main metabolic cycles (i.e., TCA, glyoxylate, and methylcitrate) are colored as well.

### Malonate utilization induces the expression of global stress response-related genes

Next, we investigated whether malonate utilization is linked to eliciting stress response in *P. aeruginosa*. Surprisingly, we identified over a hundred genes that are differentially expressed and linked to various stress responses. These can be broadly grouped as metal stress response **(Table 1)**, oxidative stress response **(Table 2)**, or carbon starvation / anaerobic stress response **(Table 3)** with a few general stress response genes that we grouped into a temperature, osmolarity, and pH stress response category **(Table 4)**.

**Table 1.**
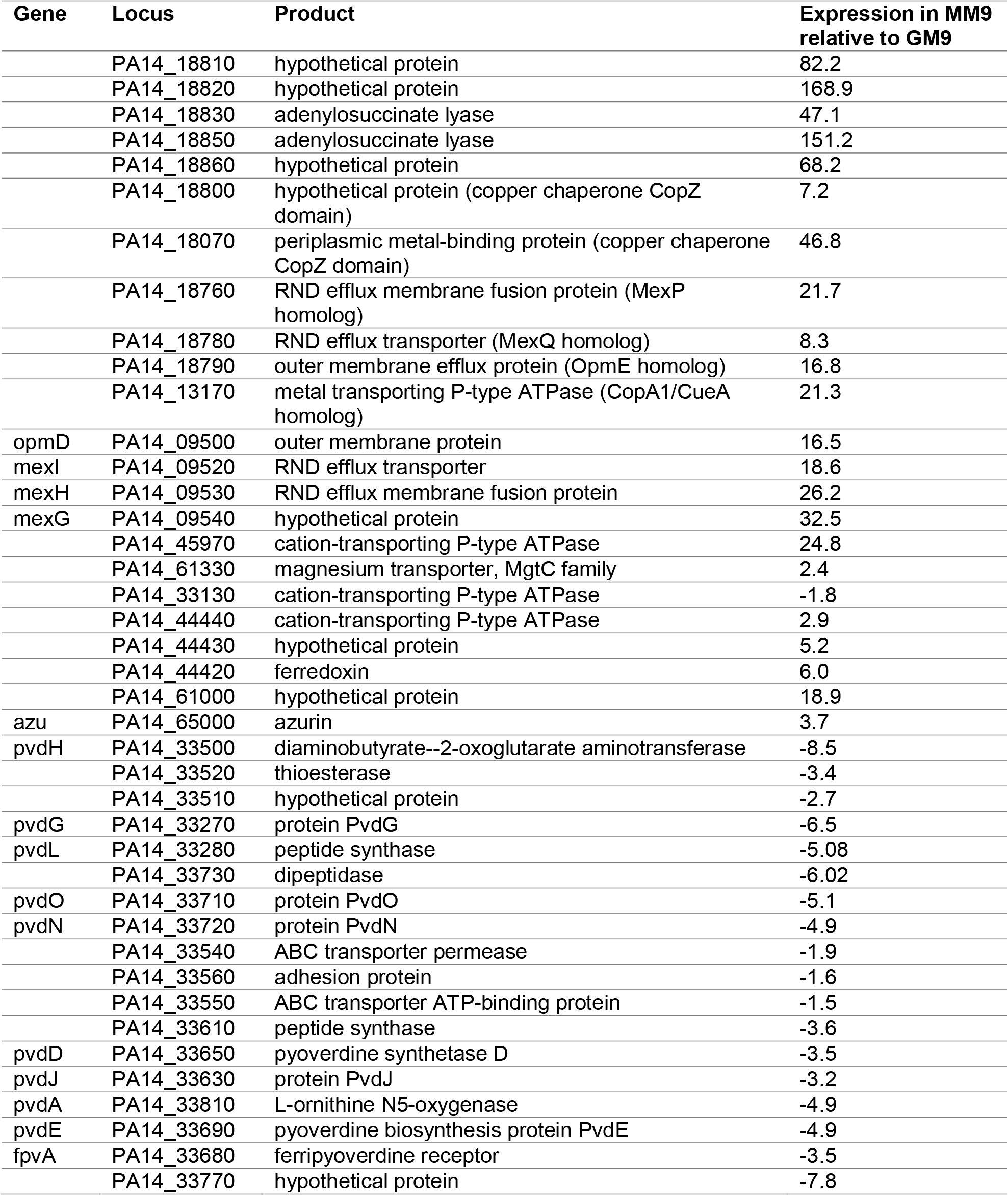
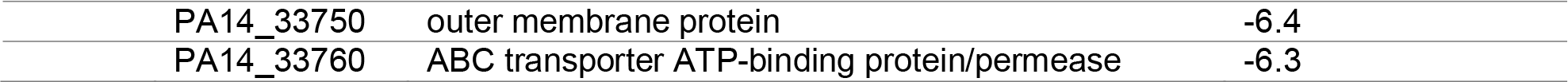
Differentially expressed genes in PA14 grown in MM9 in comparison with GM9. Listed genes are related to metal stress response.

| Gene | Locus | Product | Expression in MM9<br>relative to GM9 |
| --- | --- | --- | --- |
|  | PA14_18810 | hypothetical protein | 82.2 |
|  | PA14_18820 | hypothetical protein | 168.9 |
|  | PA14_18830 | adenylosuccinate lyase | 47.1 |
|  | PA14_18850 | adenylosuccinate lyase | 151.2 |
|  | PA14_18860 | hypothetical protein | 68.2 |
|  | PA14_18800 | hypothetical protein (copper chaperone CopZ domain) | 7.2 |
|  | PA14_18070 | periplasmic metal-binding protein (copper chaperone CopZ domain) | 46.8 |
|  | PA14_18760 | RND efflux membrane fusion protein (MexP homolog) | 21.7 |
|  | PA14_18780 | RND efflux transporter (MexQ homolog) | 8.3 |
|  | PA14_18790 | outer membrane efflux protein (OpmE homolog) | 16.8 |
|  | PA14_13170 | metal transporting P-type ATPase (CopA1/CueA homolog) | 21.3 |
| opmD | PA14_09500 | outer membrane protein | 16.5 |
| mexI | PA14_09520 | RND efflux transporter | 18.6 |
| mexH | PA14_09530 | RND efflux membrane fusion protein | 26.2 |
| mexG | PA14_09540 | hypothetical protein | 32.5 |
|  | PA14_45970 | cation-transporting P-type ATPase | 24.8 |
|  | PA14_61330 | magnesium transporter, MgtC family | 2.4 |
|  | PA14_33130 | cation-transporting P-type ATPase | -1.8 |
|  | PA14_44440 | cation-transporting P-type ATPase | 2.9 |
|  | PA14_44430 | hypothetical protein | 5.2 |
|  | PA14_44420 | ferredoxin | 6.0 |
|  | PA14_61000 | hypothetical protein | 18.9 |
| azu | PA14_65000 | azurin | 3.7 |
| pvdH | PA14_33500 | diaminobutyrate--2-oxoglutarate aminotransferase | -8.5 |
|  | PA14_33520 | thioesterase | -3.4 |
|  | PA14_33510 | hypothetical protein | -2.7 |
| pvdG | PA14_33270 | protein PvdG | -6.5 |
| pvdL | PA14_33280 | peptide synthase | -5.08 |
|  | PA14_33730 | dipeptidase | -6.02 |
| pvdO | PA14_33710 | protein PvdO | -5.1 |
| pvdN | PA14_33720 | protein PvdN | -4.9 |
|  | PA14_33540 | ABC transporter permease | -1.9 |
|  | PA14_33560 | adhesion protein | -1.6 |
|  | PA14_33550 | ABC transporter ATP-binding protein | -1.5 |
|  | PA14_33610 | peptide synthase | -3.6 |
| pvdD | PA14_33650 | pyoverdine synthetase D | -3.5 |
| pvdJ | PA14_33630 | protein PvdJ | -3.2 |
| pvdA | PA14_33810 | L-ornithine N5-oxygenase | -4.9 |
| pvdE | PA14_33690 | pyoverdine biosynthesis protein PvdE | -4.9 |
| fpvA | PA14_33680 | ferripyoverdine receptor | -3.5 |
|  | PA14_33770 | hypothetical protein | -7.8 |

|  |  |  |
| --- | --- | --- |
| PA14_33750 | outer membrane protein | -6.4 |
| PA14_33760 | ABC transporter ATP-binding protein/permease | -6.3 |

**Table 2.**
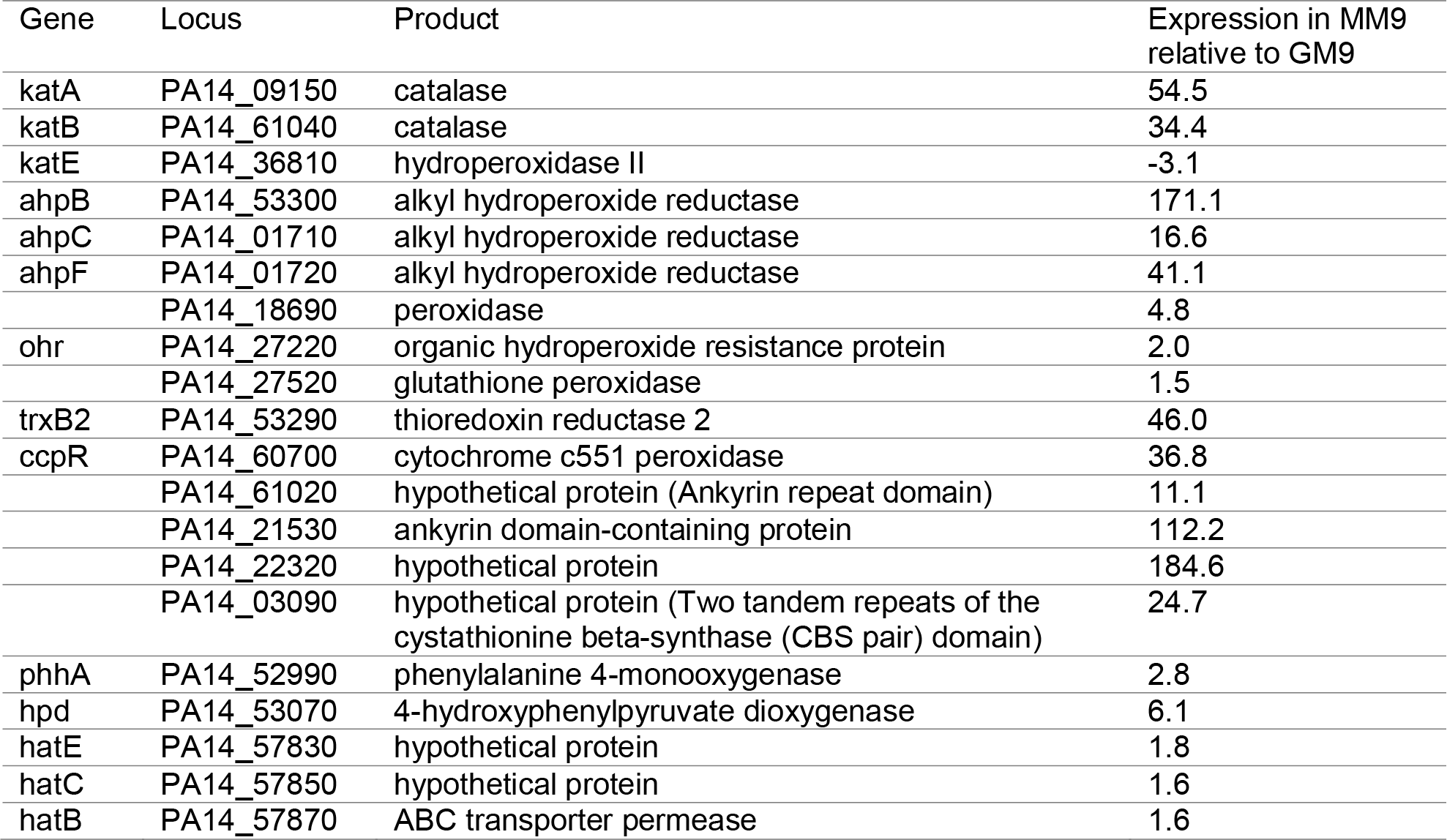
Differentially expressed genes in PA14 grown in MM9 in comparison with GM9. Listed genes are related to oxidative stress response.

**Table 3.**
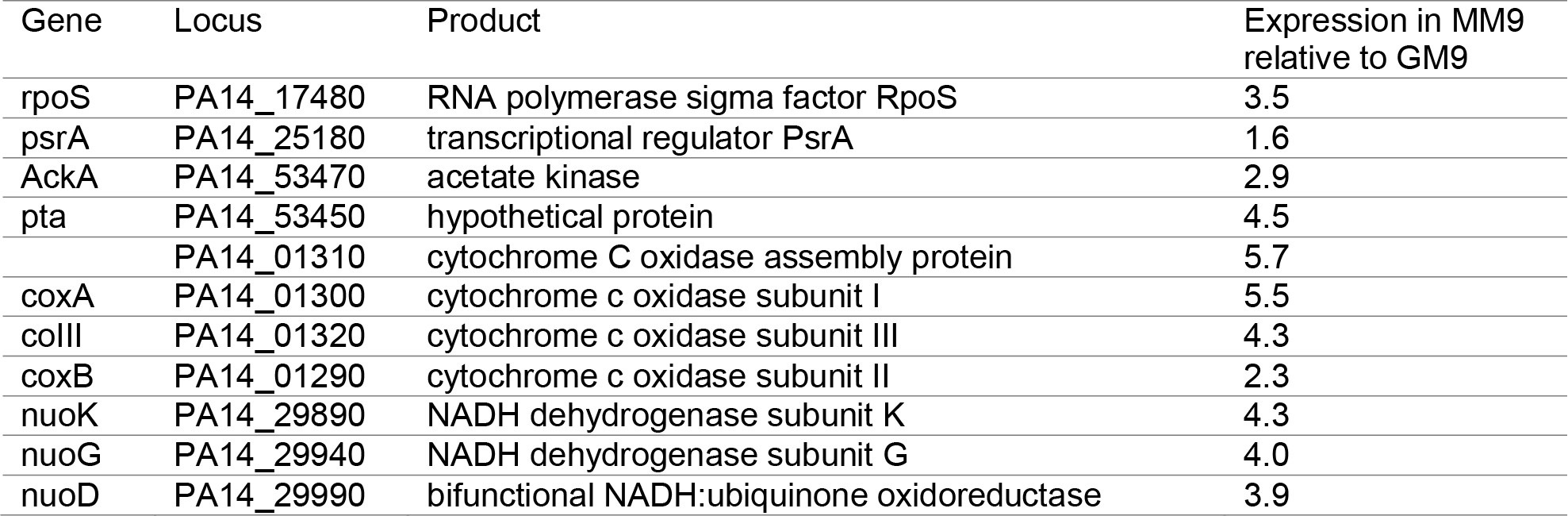
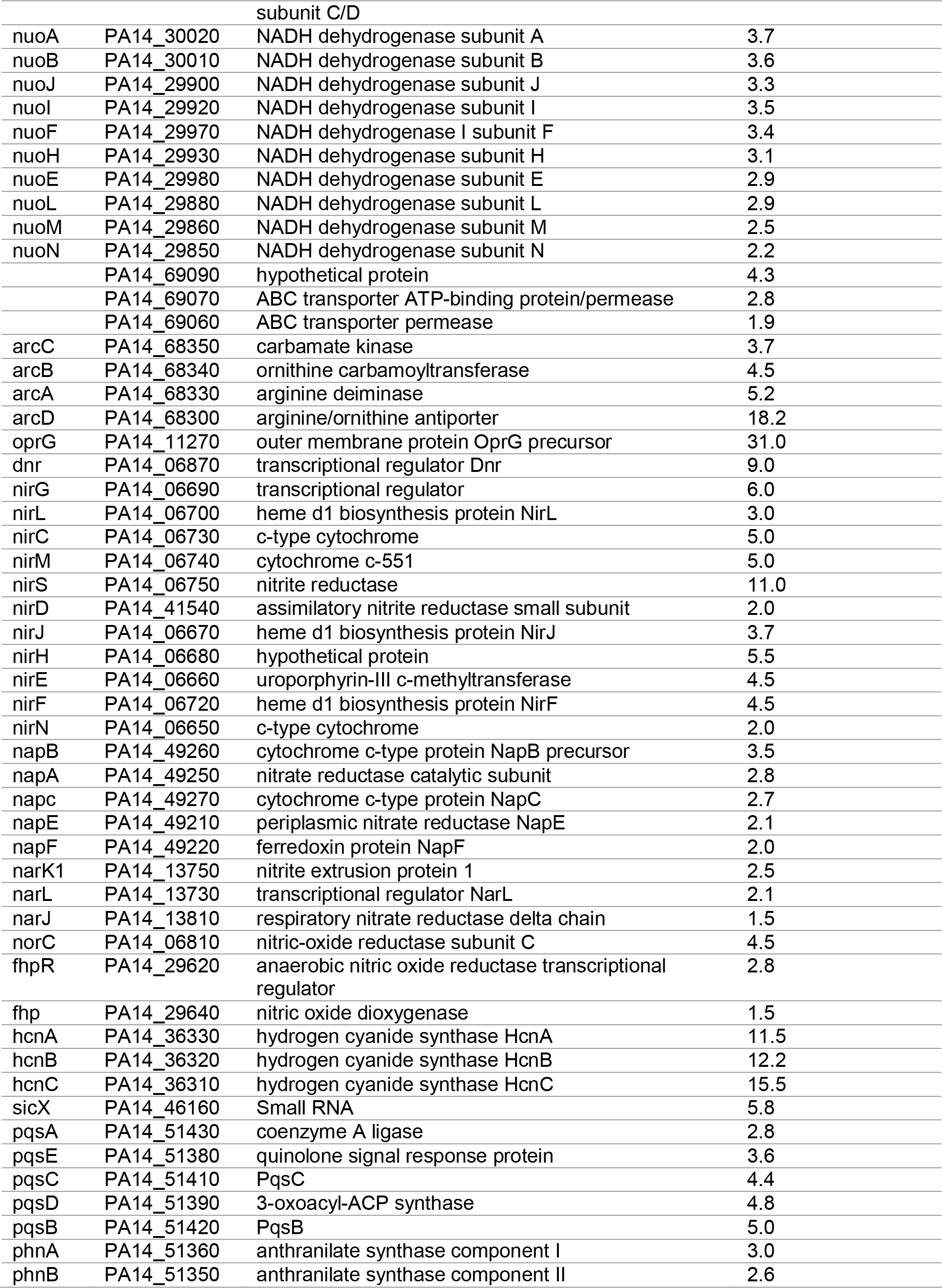
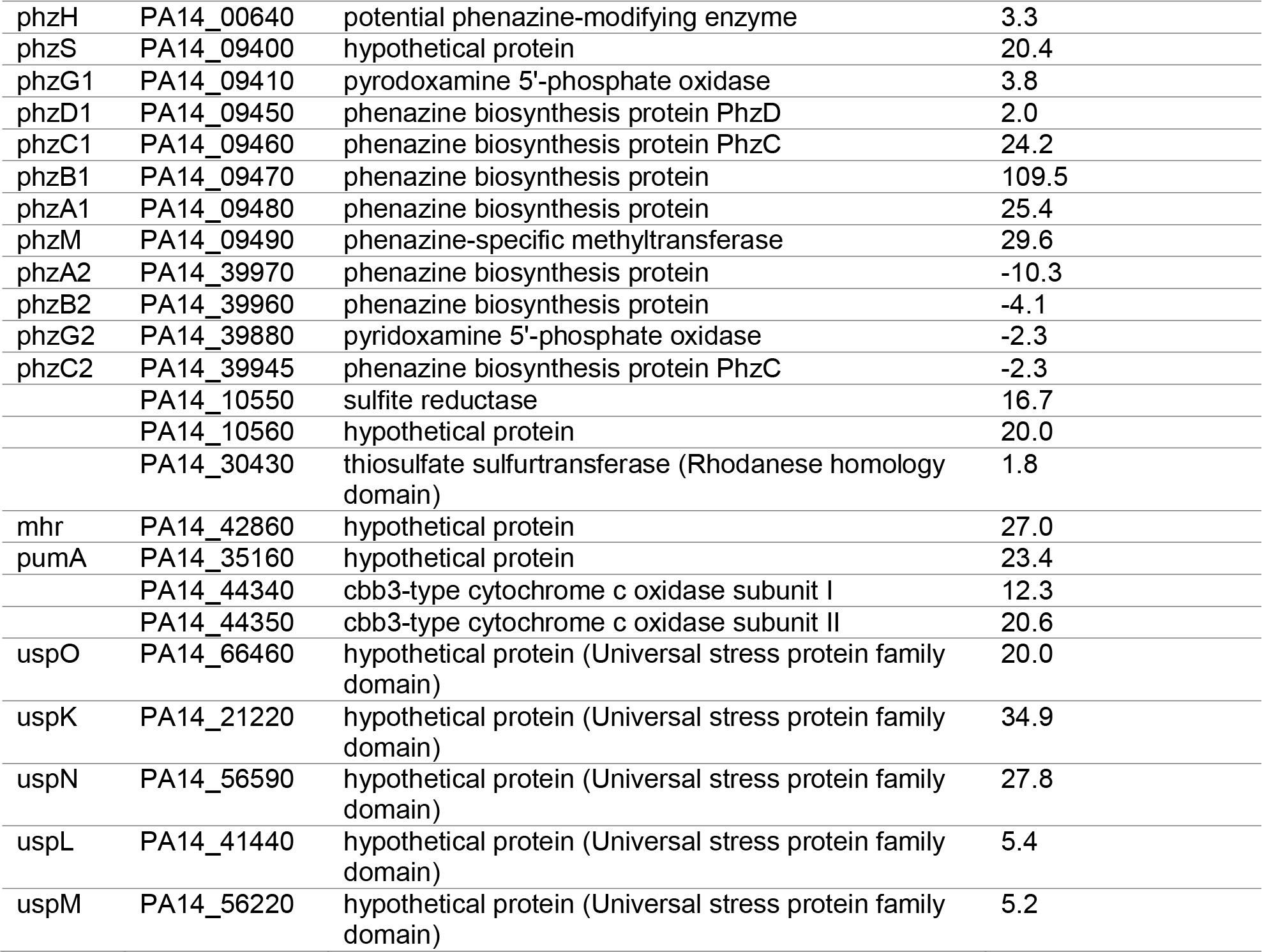
Differentially expressed genes in PA14 grown in MM9 in comparison with GM9. Listed genes are related to carbon starvation and anaerobic stress response.

| Gene | Locus | Product | Expression in MM9 relative to GM9 |
| --- | --- | --- | --- |
| rpoS | PA14_17480 | RNA polymerase sigma factor RpoS | 3.5 |
| psrA | PA14_25180 | transcriptional regulator PsrA | 1.6 |
| AckA | PA14_53470 | acetate kinase | 2.9 |
| pta | PA14_53450 | hypothetical protein | 4.5 |
|  | PA14_01310 | cytochrome C oxidase assembly protein | 5.7 |
| coxA | PA14_01300 | cytochrome c oxidase subunit I | 5.5 |
| coiII | PA14_01320 | cytochrome c oxidase subunit III | 4.3 |
| coxB | PA14_01290 | cytochrome c oxidase subunit II | 2.3 |
| nuoK | PA14_29890 | NADH dehydrogenase subunit K | 4.3 |
| nuoG | PA14_29940 | NADH dehydrogenase subunit G | 4.0 |
| nuoD | PA14_29990 | bifunctional NADH:ubiquinone oxidoreductase subunit C/D | 3.9 |
| nuoA | PA14_30020 | NADH dehydrogenase subunit A | 3.7 |
| nuoB | PA14_30010 | NADH dehydrogenase subunit B | 3.6 |
| nuoJ | PA14_29900 | NADH dehydrogenase subunit J | 3.3 |
| nuoI | PA14_29920 | NADH dehydrogenase subunit I | 3.5 |
| nuoF | PA14_29970 | NADH dehydrogenase I subunit F | 3.4 |
| nuoH | PA14_29930 | NADH dehydrogenase subunit H | 3.1 |
| nuoE | PA14_29980 | NADH dehydrogenase subunit E | 2.9 |
| nuoL | PA14_29880 | NADH dehydrogenase subunit L | 2.9 |
| nuoM | PA14_29860 | NADH dehydrogenase subunit M | 2.5 |
| nuoN | PA14_29850 | NADH dehydrogenase subunit N | 2.2 |
|  | PA14_69090 | hypothetical protein | 4.3 |
|  | PA14_69070 | ABC transporter ATP-binding protein/permease | 2.8 |
|  | PA14_69060 | ABC transporter permease | 1.9 |
| arcC | PA14_68350 | carbamate kinase | 3.7 |
| arcB | PA14_68340 | ornithine carbamoyltransferase | 4.5 |
| arcA | PA14_68330 | arginine deiminase | 5.2 |
| arcD | PA14_68300 | arginine/ornithine antiporter | 18.2 |
| oprG | PA14_11270 | outer membrane protein OprG precursor | 31.0 |
| dnr | PA14_06870 | transcriptional regulator Dnr | 9.0 |
| nirG | PA14_06690 | transcriptional regulator | 6.0 |
| nirL | PA14_06700 | heme d1 biosynthesis protein NirL | 3.0 |
| nirC | PA14_06730 | c-type cytochrome | 5.0 |
| nirM | PA14_06740 | cytochrome c-551 | 5.0 |
| nirS | PA14_06750 | nitrite reductase | 11.0 |
| nirD | PA14_41540 | assimilatory nitrite reductase small subunit | 2.0 |
| nirJ | PA14_06670 | heme d1 biosynthesis protein NirJ | 3.7 |
| nirH | PA14_06680 | hypothetical protein | 5.5 |
| nirE | PA14_06660 | uroporphyrin-III c-methyltransferase | 4.5 |
| nirF | PA14_06720 | heme d1 biosynthesis protein NirF | 4.5 |
| nirN | PA14_06650 | c-type cytochrome | 2.0 |
| napB | PA14_49260 | cytochrome c-type protein NapB precursor | 3.5 |
| napA | PA14_49250 | nitrate reductase catalytic subunit | 2.8 |
| napC | PA14_49270 | cytochrome c-type protein NapC | 2.7 |
| napE | PA14_49210 | periplasmic nitrate reductase NapE | 2.1 |
| napF | PA14_49220 | ferredoxin protein NapF | 2.0 |
| narK1 | PA14_13750 | nitrite extrusion protein 1 | 2.5 |
| narL | PA14_13730 | transcriptional regulator NarL | 2.1 |
| narJ | PA14_13810 | respiratory nitrate reductase delta chain | 1.5 |
| norC | PA14_06810 | nitric-oxide reductase subunit C | 4.5 |
| fhpR | PA14_29620 | anaerobic nitric oxide reductase transcriptional regulator | 2.8 |
| fhp | PA14_29640 | nitric oxide dioxygenase | 1.5 |
| hcnA | PA14_36330 | hydrogen cyanide synthase HcnA | 11.5 |
| hcnB | PA14_36320 | hydrogen cyanide synthase HcnB | 12.2 |
| hcnC | PA14_36310 | hydrogen cyanide synthase HcnC | 15.5 |
| sicX | PA14_46160 | Small RNA | 5.8 |
| pqsA | PA14_51430 | coenzyme A ligase | 2.8 |
| pqsE | PA14_51380 | quinolone signal response protein | 3.6 |
| pqsC | PA14_51410 | PqsC | 4.4 |
| pqsD | PA14_51390 | 3-oxoacyl-ACP synthase | 4.8 |
| pqsB | PA14_51420 | PqsB | 5.0 |
| phnA | PA14_51360 | anthranilate synthase component I | 3.0 |
| phnB | PA14_51350 | anthranilate synthase component II | 2.6 |
| phzH | PA14_00640 | potential phenazine-modifying enzyme | 3.3 |
| phzS | PA14_09400 | hypothetical protein | 20.4 |
| phzG1 | PA14_09410 | pyrodoxamine 5'-phosphate oxidase | 3.8 |
| phzD1 | PA14_09450 | phenazine biosynthesis protein PhzD | 2.0 |
| phzC1 | PA14_09460 | phenazine biosynthesis protein PhzC | 24.2 |
| phzB1 | PA14_09470 | phenazine biosynthesis protein | 109.5 |
| phzA1 | PA14_09480 | phenazine biosynthesis protein | 25.4 |
| phzM | PA14_09490 | phenazine-specific methyltransferase | 29.6 |
| phzA2 | PA14_39970 | phenazine biosynthesis protein | -10.3 |
| phzB2 | PA14_39960 | phenazine biosynthesis protein | -4.1 |
| phzG2 | PA14_39880 | pyridoxamine 5'-phosphate oxidase | -2.3 |
| phzC2 | PA14_39945 | phenazine biosynthesis protein PhzC | -2.3 |
|  | PA14_10550 | sulfite reductase | 16.7 |
|  | PA14_10560 | hypothetical protein | 20.0 |
|  | PA14_30430 | thiosulfate sulfurtransferase (Rhodanese homology domain) | 1.8 |
| mhr | PA14_42860 | hypothetical protein | 27.0 |
| pumA | PA14_35160 | hypothetical protein | 23.4 |
|  | PA14_44340 | cbb3-type cytochrome c oxidase subunit I | 12.3 |
|  | PA14_44350 | cbb3-type cytochrome c oxidase subunit II | 20.6 |
| uspO | PA14_66460 | hypothetical protein (Universal stress protein family domain) | 20.0 |
| uspK | PA14_21220 | hypothetical protein (Universal stress protein family domain) | 34.9 |
| uspN | PA14_56590 | hypothetical protein (Universal stress protein family domain) | 27.8 |
| uspL | PA14_41440 | hypothetical protein (Universal stress protein family domain) | 5.4 |
| uspM | PA14_56220 | hypothetical protein (Universal stress protein family domain) | 5.2 |

**Table 4.**
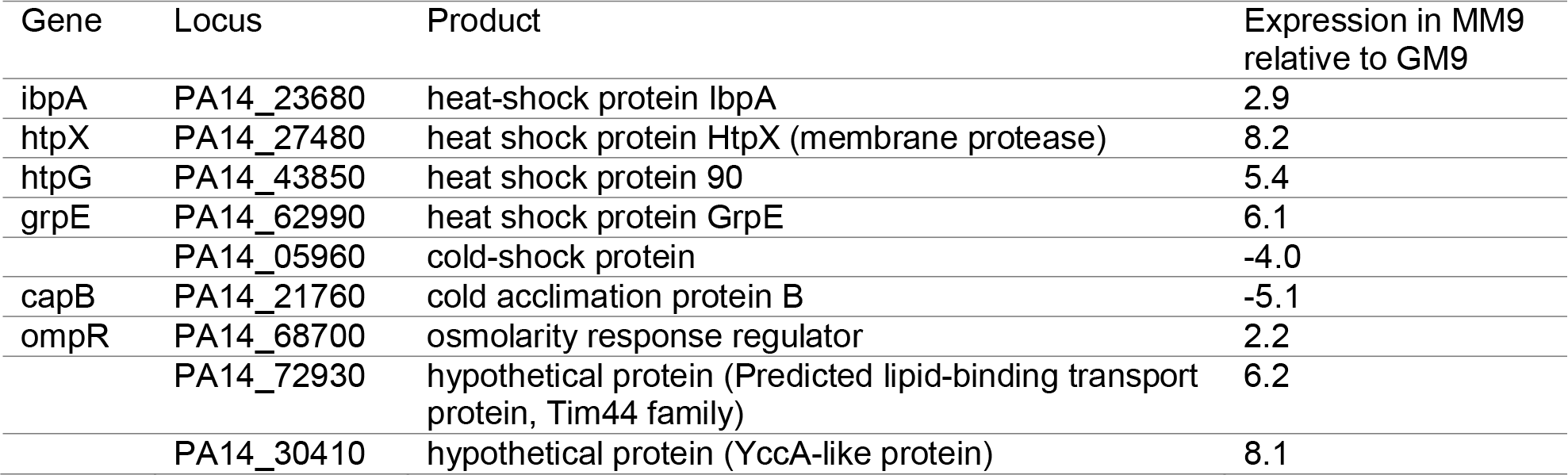
Differentially expressed genes in PA14 grown in MM9 in comparison with GM9. Listed genes are related to temperature, osmolarity, and pH stress responses.

### Metals and oxidative stress response

Our transcriptomic analysis revealed that malonate utilization relative to glycerol utilization induces genes that are associated with copper toxicity response in PA14 **(Table 1)** [19]. Genes representing the copper resistance regulon include the operon consisting of the five genes *PA14_18810*-*18860* that was upregulated with an average of 103.5 times higher in MM9 than in GM9. Two copper-related genes, *PA14_18800* and PA14_18070, were induced 7.2 and 46.8 times, respectively. These genes code for proteins with a copper chaperone CopZ domain. Three genes (i.e., *PA14_18760*-*18790*) coding for a homolog of the resistance-nodulation-division (RND) efflux pump MexPQ-OpmE were also induced with an average of 15.6 times. This efflux pump was observed to be activated by copper [20]. *PA14_13170*, a *copA1*/*cueA* homolog (a P-type ATPase), was induced 21.3 times. These copper resistance-regulated genes are known to be controlled by the transcriptional regulator CueR, and mutations in any of these genes result in a higher sensitivity to copper [19]. Surprisingly, this gene’s homolog (*PA14_63170*) was not differentially expressed (P-value = 0.2). Several other genes related to metal stress response were differentially regulated with malonate utilization.

For example, the four genes coding for the RND efflux pump, MexGHI-OpmD, were upregulated with an average of 23.5 times. Other genes included those coding for cation-transporting P-type ATPases **(Table 1)**. Lastly, *PA14_65000* that codes for azurin was upregulated 3.7 times. We also observed that the genes involved in pyoverdine biosynthesis and uptake were downregulated in MM9, suggesting that the cell is shutting down its metal import systems **(Table 1)**. The *pvdD* gene was downregulated 3 times, while the pyoverdine biosynthesis protein PvdE was downregulated 5 times. These results suggest that malonate utilization results in metal stress response due to either the accumulation of metals accompanied by malonate utilization or an alteration in the regulatory network controlling metals sensing— independent of metal concentration.

Because oxidative stress is strongly linked to metal stress, we aimed next to investigate if oxidative stress-related genes were also differentially regulated by malonate utilization [21, 22]. Indeed, over a dozen genes involved in oxidative response were upregulated **(Table 2)**. Catalase-coding genes (*katA* and *katB*) were upregulated 54.5 and 34.4 times, but *katE* was downregulated 3.1 times. *PA14_61020* is a gene in the same operon as *katB* was found to be upregulated 11.1 times. Its gene product has an ankyrin repeat domain, which plays a role in signal transduction [23]. *PA14_21530* is another gene harboring ankyrin repeat domain in its protein product that was significantly upregulated 112.2 times. Interestingly, the expression of this gene is positively regulated by MvfR [24] and associated with oxidative stress adaptation [25]. *PA14_22320* is another gene that was significantly upregulated 184.6 times. It is possibly controlled by MvfR [24], and its expression is induced by hydroxyl radicals [26]. *PA14_03090* was another gene that was upregulated, and evidence suggests that it is required for hydrogen peroxide resistance [25]. Three alkyl hydroperoxide reductase- coding genes (*ahpB*, *ahpC*, and *ahpF*) were upregulated with an average of 76.3 times—among them, *ahpB* differential expression was the highest, 171.1 times. Five peroxidase-coding genes were upregulated such as glutathione peroxidase, thioredoxin reductase 2, and cytochrome c551 peroxidase. These results are consistent with our previous finding where we observed an increase in the catalase activity of PA14 grown in MM9 [14].

Pyomelanin is a pigment produced by *P. aeruginosa*, and one of its main functions is to provide protection against oxidative stress [27]. Our results suggest that malonate utilization is associated with its upregulation **(Table 2)**. Genes *PA14_52990* and *PA14_53070*, which are involved in the biosynthesis of homogentisate (the precursor of pyomelanin) from chorismate were upregulated. Moreover, genes (*PA14_57830*, *PA14_57850*, and *PA14_57870*) involved in the transport of homogentisate [28] and possibly the extracellular accumulation of pyomelanin were upregulated as well. Overall, these results suggest that malonate utilization in *P. aeruginosa* is accompanied by oxidative stress response.

### Carbon starvation and anaerobic stress response

Just as metal stress and oxidative stress are inseparably linked, carbon starvation and anaerobic stress response share many facets. Transcriptomic analysis of *P. aeruginosa* during malonate utilization reveals that *P. aeruginosa* is undergoing carbon starvation and anaerobic stress response **(Table 3)**. The expression of *rpoS*, which encodes a sigma-factor known to be associated with carbon or oxygen limitation [29], was upregulated 3.5 times. The expression of *psrA*, an *rpoS* transcriptional regulator [30], was also upregulated 1.6 times. The *aa*_3_-type cytochrome *c* oxidase, encoded by the *coxBA-PA14_01310-coIII* gene cluster and induced by RpoS upon carbon limitation [3], was also found to be upregulated an average of 4.4 times.

All 13 genes (*nuoN*, *nuoM*, *nuoL*, *nuoK*, *nuoJ*, *nuoI*, *nuoH*, *nuoG*, *nuoF*, *nuoE*, *nuoD*, *nuoB*, and *nuoA*) constituting the *nuo* operon were upregulated on average 3.3 times **(Table 3)**. This operon codes for the NADH dehydrogenase (NADH: quinone oxidoreductase). This enzyme translocates protons, oxidizes NADH to NAD^+^ [31], and is required for anaerobic growth [32]. Furthermore, this system is important for *P. aeruginosa* virulence and provides resistance towards aminoglycoside antibiotics [33].

We also identified a putative ABC transporter encoded by *PA14_69090*, *PA14_69070*, and *PA14_69060* that was upregulated ∼3 times. A previous study suggests that this ABC transporter is also required during oxygen limitation [30]. The expression of the outer-membrane protein encoded by *oprG*, was upregulated 31 times. OprG is a specific transporter of hydrophobic molecules and its expression is induced under anaerobic conditions [34].

Five genes encoding proteins with the universal stress protein family domains were upregulated **(Table 3)**. These included *uspO*, *uspK*, *uspN*, *uspL*, and *uspM* with an average increase of 18.7 times. These genes play a role in response to different stressors—more importantly, anaerobic stress conditions or oxygen limitation [35]. All genes of *arcDABC* operon, which is responsible for arginine fermentation, were upregulated. The expression of *uspK*, *uspN* and *arcDABC* was reported to be induced in biofilms or anaerobic environments [36]. Two genes (*acka* and *pta*) involved in pyruvate fermentation and anaerobic metabolism [37] were also upregulated. Moreover, the *cbb*_3_-type cytochrome c oxidase, encoded by *PA14_44340* and *PA14_44350*, was also upregulated. This system is induced under an oxygen limited environment [38]. Lastly, another recently identified gene that is associated with survival under oxygen-low condition, *mhr,* was found to be upregulated almost 27 times [39].

The *dnr* gene (*PA14_06870*)-encoding the transcriptional regulator Dnr was upregulated 9 times **(Table 3)**. Dnr senses nitric oxide [40] and regulates the denitrification gene [41]. During the denitrification process, nitrate is used as an alternative terminal electron acceptor and reduced to nitrogen [42]. Our results showed that 20 genes that are part of the denitrification operons (*nar*, *nir*, *nap*, *nor*, and *nos*) were induced. Our finding of increased expression of many *nos, nor*, *nir*, *and dnr* genes by cells suggests that *P. aeruginosa* is exposed to anaerobic conditions when grown in malonate as a sole carbon source. The *fhp* gene, and its transcriptional regulator *fhpR*, were upregulated as well. This flavohemoglobin system is nitric oxide-responsive and impairs the dispersal response to nitric oxide [43]. These results suggest that nitric oxide is produced during malonate utilization, thus eliciting an anaerobic (low oxygen) stress response. This was supported by the upregulation in the three genes constituting *hcn* operon that is responsible for the synthesis of hydrogen cyanide, which is induced under hypoxic environment [44]. Moreover, the recently characterized small RNA, *sicX*, encoded by PA14_46160 was upregulated ∼6 times. It is worth noting that *sicX* was found to be induced by low oxygen and is important in the transition between chronic and acute infection stages in *P. aeruginosa* [45].

Eight genes involved in the biosynthesis of pyocyanin were significantly upregulated **(Table 3)**. For example, *phzB_1_*, one of the main biosynthetic operons *phz_1_*, was upregulated 109.5 times. But four genes of the redundant operon *phz_2_*were downregulated. This differential regulation of these biosynthetic operons is worth addressing; however, it is beyond the scope of this study. This possible increase in pyocyanin fits with our hypothesis, as pyocyanin is important for *P. aeruginosa* to maintain redox homeostasis and survive in biofilms under oxygen-depleted environments [3, 46]. The upregulation of the phenazine biosynthesis genes is further supported by the upregulation in biosynthetic genes (*phnAB* and *pqs* operon) of *Pseudomonas* quinolone signal (PQS), which tightly regulates pyocyanin production. Also, our data is consistent with our prior report that showed that growth in MM9 impacts pyocyanin production and quorum-sensing regulated pathways [14]. *P. aeruginosa* has multiple resistance mechanisms to protect itself from the toxic effects of pyocyanin and maintain redox homeostasis [47]. This was observed through the upregulation of expression of genes encoding the monooxygenase PumA and RNA efflux pump MexGHI-OpmG. One operon consisting of two genes associated with cyanide resistance (i.e., *PA14_10550* and *PA14_10560*) was upregulated [48, 49].

Besides this operon, a thiosulfate sulfur-transferase encoded by *PA14_30430* was upregulated. Although this is not experimentally confirmed yet, we hypothesize that the thiosulfate sulfur-transferase (encoded by *PA14_30430*), which carries repeated rhodanese homology domains, detoxifies cyanide to sulfite and thiocyanate. Then, the sulfite reductase (encoded by *PA14_10550*) reduces sulfite to sulfide. Whether the hypothetical protein encoded by *PA14_10560* is possibly involved is yet to be determined. Overall, the results suggest that malonate utilization is tightly linked with anaerobic respiration and associated with carbon starvation stress response.

### Temperature, osmolarity, and pH stress response

Despite the fact that all cultures were incubated at 37°C, malonate utilization was associated with heat-shock response on the transcriptomic level **(Table 4)**. This was accompanied by an increase in the expression of four heat-shock genes encoding for proteins (IbpA, HtpX, HtpG, and GrpE) with an average of 5.6 times and decrease in the expression of two cold-shock proteins with an average of 4.6 times. Our results are consistent with the findings of other groups that indicated heat-shock responses can be induced by conditions that did not alter temperature such as alkaline pH or carbon starvation [50–52].

Finally, we observed the upregulation of certain genes involved in osmolarity and pH fluctuations **(Table 4)**. For example, the expression of the transcriptional regulator OmpR was significantly upregulated. Among its regulated genes are *htpX* (membrane protease), *PA14_72930* (predicted lipid-binding transport protein, Tim44 family), and *PA14_30410* (YccA-like protein). All of *ompR*, *htpX*, *PA14_72930*, and *PA14_30410* protect *P. aeruginosa* against osmolarity and pH stressors [53]. The role of these genes involves maintaining membrane integrity. Because of this critical role, the function of those genes involves the protection against aminoglycoside antibiotics [53]. Overall, these results suggest that malonate utilization induces a global stress response in *P. aeruginosa* that spans various physical and physiological stressors. However, it is not clear what these adaptations are responsive to exactly, which we aimed to address next.

### Malonate utilization modulates the intracellular accumulation of certain metals

Because the transcriptomic analysis revealed a striking metal stress response, we hypothesized that malonate utilization results in the accumulation of metal ions within the bacterial cells. To determine the effect of malonate or glycerol as sole carbon sources on the accumulation of various metal ions within *P. aeruginosa* cells, we used inductively coupled plasma mass spectrometry (ICP-MS) to detect the level of metal ions. Because chemicals may carry metal impurities, we used Chelex-treated M9 and Chelex-treated carbon sources. Chelex is a chelating agent that binds polyvalent metal ions, thus this treatment removed the possibility that metal changes in the different treatments could be driven by trace metal contaminates in the different carbon sources. This procedure eliminated several metals essential for growth, therefore, to complement those metals, we added a cocktail of these essential metal ions to the prepared media (i.e., Mg 1mM, Mn 25 µM, Ca 100 µM, Fe 5 µM, Zn 25 µM, Co 0.1 µM, and Cu 0.1 µM). After overnight incubation of *P. aeruginosa* in Chelex-treated GM9 and MM9, cells were collected and then prepared using an established protocol following slight modifications [54]. Using ICP-MS analysis of the overnight cultures we measured the concentration of 17 metal ions **(Figure 2)**. While most of the measured metal ions were similarly abundant in cells obtained from GM9 and MM9, two metal ions (i.e., Al and Sr) showed a statistically significant difference between MM9 and GM9. Specifically, Al and Sr were ∼4 and ∼28 times, respectively, more abundant intracellularly in MM9 versus GM9 **(Figure 2)**. The mechanism by which malonate utilization induces the accumulation of metal ions is not clear yet. One possible explanation is that due to the lower level of pyoverdine in MM9 and its ability to decrease metal accumulation, this may be responsible for the accumulation of AI intracellularly in *P. aeruginosa* grown in MM9 [55, 56].

**Figure 2.**
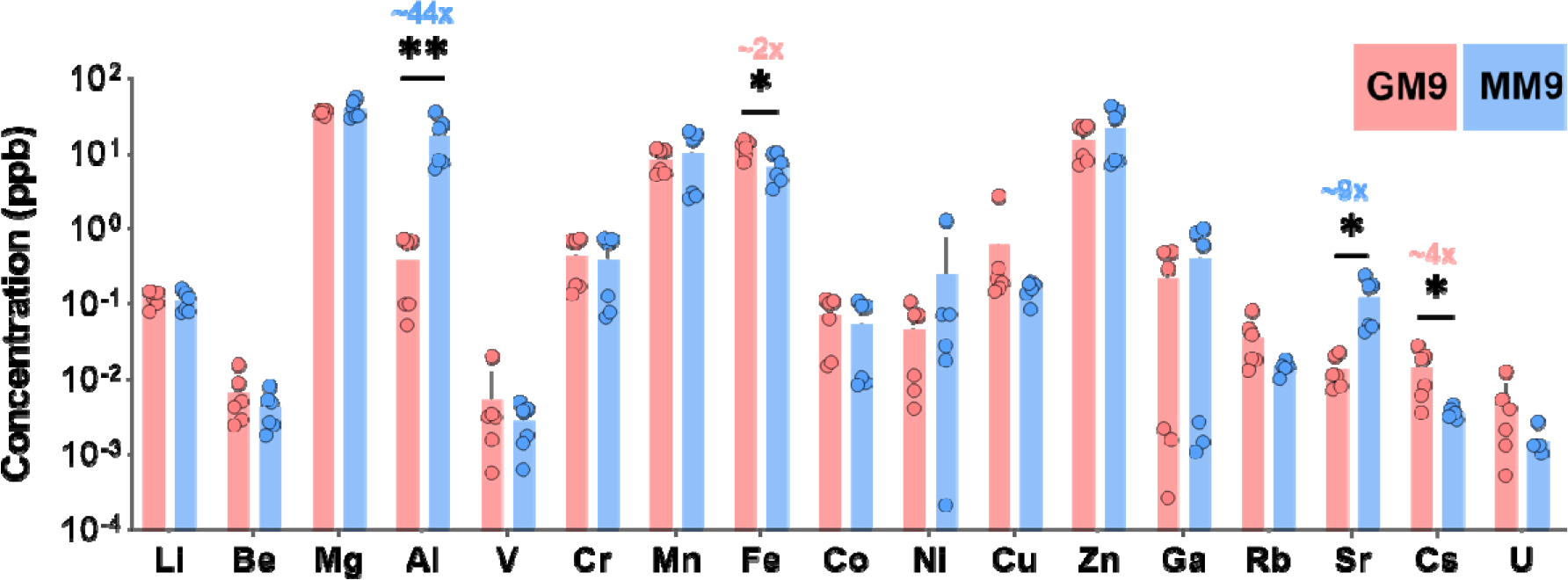
Effect of malonate utilization on accumulation of metal ions in *P. aeruginosa.* (A) Accumulation level of all measured metal ions using ICP-MS. **(B)** Ratio of Al and Sr accumulation level in MM9 in comparison to GM9. PA14 was grown in Chelex-treated malonate and glycerol supplemented Chelex-treated M9. Samples were processed for ICP-MS analysis as mentioned in the Experimental Procedures. Samples were normalized based on protein content, then analyzed for the accumulation of the metal ions using ICP-MS. Error bars represent standard deviations of the results from triplicate samples. Unpaired t-test (two-tailed) was used to measure statistical significance. * P < 0.05 and ** P < 0.01. Concentration of metal ions was measured in parts per billion (ppb). This experiment was repeated three times on three different days and repeated once using the previously used carbon source concentrations and once using equimolar concentrations.

Because we observed an upregulation in all known regulons related to copper stress response, we expected to find Cu to be accumulated in MM9. Surprisingly, this was not the case **(Figure 2)**. One possible explanation is that while Cu gets accumulated at first, *P. aeruginosa* then adapts to get rid of excess Cu through the production of a secreted copper-containing small molecule named fluopsin C [57]. This hypothesis is strengthened by the prominent upregulation (over 100 times higher in MM9 vs. GM9) of the biosynthetic operon consisting of five genes *PA14_18810*-*18860* that is responsible for fluopsin C biosynthesis.

Motivated by the phenotype of *P. aeruginosa*, in which its malonate utilization induces the expression of biosynthetic genes of fluopsin C and pyocyanin, and knowing both of these molecules act as antibiotics against *Staphylococcus aureus* [57, 58], we hypothesized that this could benefit *P. aeruginosa* in its competition against other bacteria it may encounter in the lung environment. We further hypothesized that the survival of *S. aureus* will decrease when co-cultured with *P. aeruginosa* in MM9, but not in GM9. To test this hypothesis, we used *S. aureus* strain JE2, an established methicillin-resistant lab strain [59], to study its survival in the presence of *P. aeruginosa* in MM9 and GM9. A co-culture of *P. aeruginosa* and *S. aureus* in a 1:1 ratio was grown in both MM9 and GM9 overnight. Then, cells were serially diluted and plated on selective media. First, we confirmed that no difference was observed in the growth of *P. aeruginosa* when co-cultured with *S. aureus* **(Supplementary** Figure 4A). In contrast, we observed a significant reduction in the number of *S. aureus* cells when co-cultured with *P. aeruginosa* in MM9, while no such difference was observed when co-cultured with *P. aeruginosa* in GM9 **(Supplementary** Figure 4B). While the higher levels of pyocyanin and possibly fluopsin C in MM9 as compared to GM9 may explain the increased death rate of *S. aureus* in MM9, other virulence or metabolic factors may contribute to this phenotype as well.

### Genetic requirements for *P. aeruginosa* to grow using malonate as a sole carbon source

The observed upregulation in many stress-related genes raises an interesting question: Are these stress-related genes essential for the growth of *P. aeruginosa* in MM9 to cope with the stress induced by malonate utilization? To address this question, we performed a high-throughput screening on a commercially available transposon mutant library of *P. aeruginosa* UCBPP-PA14 containing over 5,500 unique mutants [60]. We grew wild-type PA14 and the mutants in GM9 and MM9 and measured bacterial growth using optical density (OD_600_). We observed a variable number of preliminary candidate mutants showing no growth in either MM9, GM9, or both **(Supplementary Table 1)**. Upon the completion of the primary screening, we selected the hits of interest that overlapped with our transcriptomics dataset for further testing.

We selected these mutants along with the wild-type (WT) PA14 to re-assess their growth in MM9 and GM9 **(Figure 3A and 3B).** While some mutants showed a statistically significant difference in their growth in comparison to that of WT, only Δ*PA14_27480* showed remarkable defective growth in both MM9 and GM9. The expression of *PA14_27480* was upregulated 8 times in MM9 vs. GM9 (**Table 4**). *PA14_27480* codes for the heat shock protein HtpX. HtpX and other heat-shock proteases allow *P. aeruginosa* to cope with protein misfolding stress induced by high temperature, carbon starvation, or alkaline pH thus promoting its survival [52]. Eight mutants showed differential growth in MM9 or GM9, four of which showed more defective growth in MM9 rather than GM9 **(Figure 3C)**. Δ*PA14_18850* caught our attention because its expression was ∼150 times higher in MM9 vs. GM9. Furthermore, *PA14_18850* codes for FlcB, which catalyzes the first step in fluopsin C biosynthesis.

**Figure 3.**
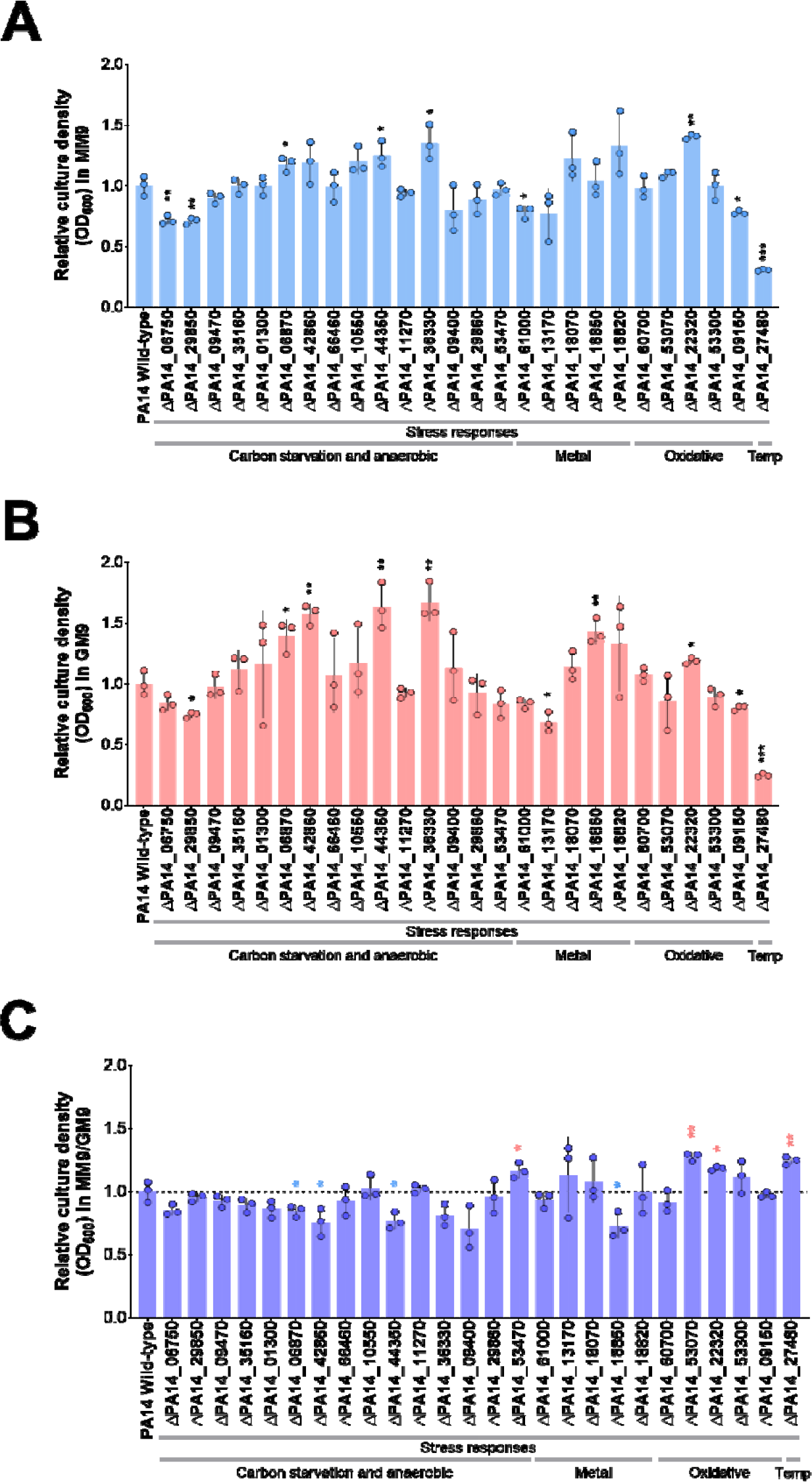
The growth of different *P. aeruginosa* strains that carry mutations in genes involved in stress adaptation. (A) Growth in strains in MM9**. (B)** Growth in strains in GM9**. (C)** MM9/GM9 culture density ratio. Bacteria were grown for 24 hours before measuring OD_600_. The mutants were run in three replicates, on three independent days. Bars represent the mean of three biological replicates. Error bars represent the standard deviation of the replicates. Data points were normalized to PA14 wild-type mean. Unpaired t-test (two-tailed) was used to measure statistical significance. * P < 0.05, ** P < 0.01, and *** P < 0.001. Statistical significance indicates statistical difference between the growth of the mutant vs. that of wild-type PA14. Temp: Temperature.

Because fluopsin C plays a role in copper detoxification, this suggests that this functional adaptation is important for *P. aeruginosa* to cope with metal stress response in MM9.

### Conservation of malonate associated phenotypes across lab and clinical strains

Thus far, we have shown that malonate utilization by *P. aeruginosa* is associated with unique properties evident at both transcriptional and phenotypic levels, e.g., pyocyanin overproduction and cell aggregation [14]. However, this is based on only one lab strain, i.e., PA14, and it is unclear if these observations are conserved in other *P. aeruginosa* strains. To this end, our next goal was to examine whether the malonate utilization-associated phenotypes are consistent across other *P. aeruginosa* lab strains **(Table 5)**. We used the widely used lab strains PAO1 [61] and PAK [62] and examined their culture growth, pyocyanin level, and cell aggregation in MM9 and GM9 [14]. While the growth of PAK strain was comparable to that of PA14 in MM9, PAO1 showed lower growth density in MM9 when compared to that of PA14 **(Figure 4A)**. Next, we quantified pyocyanin concentration produced by all strains using an established protocol [63]. Both PAO1 and PAK produced a very low level of pyocyanin in both GM9 and MM9, which was strikingly different than that of PA14 in MM9 **(Figure 4B)**. Lastly, cell aggregation phenotype, which we have observed before with PA14 grown in MM9 [14], was reproducible in all strains grown in MM9 but not in GM9 **(Figure 4C)**. Overall, these results suggest that while some phenotypes associated with malonate utilization are conserved across lab strains, others are unique to each lab strain.

**Figure 4.**
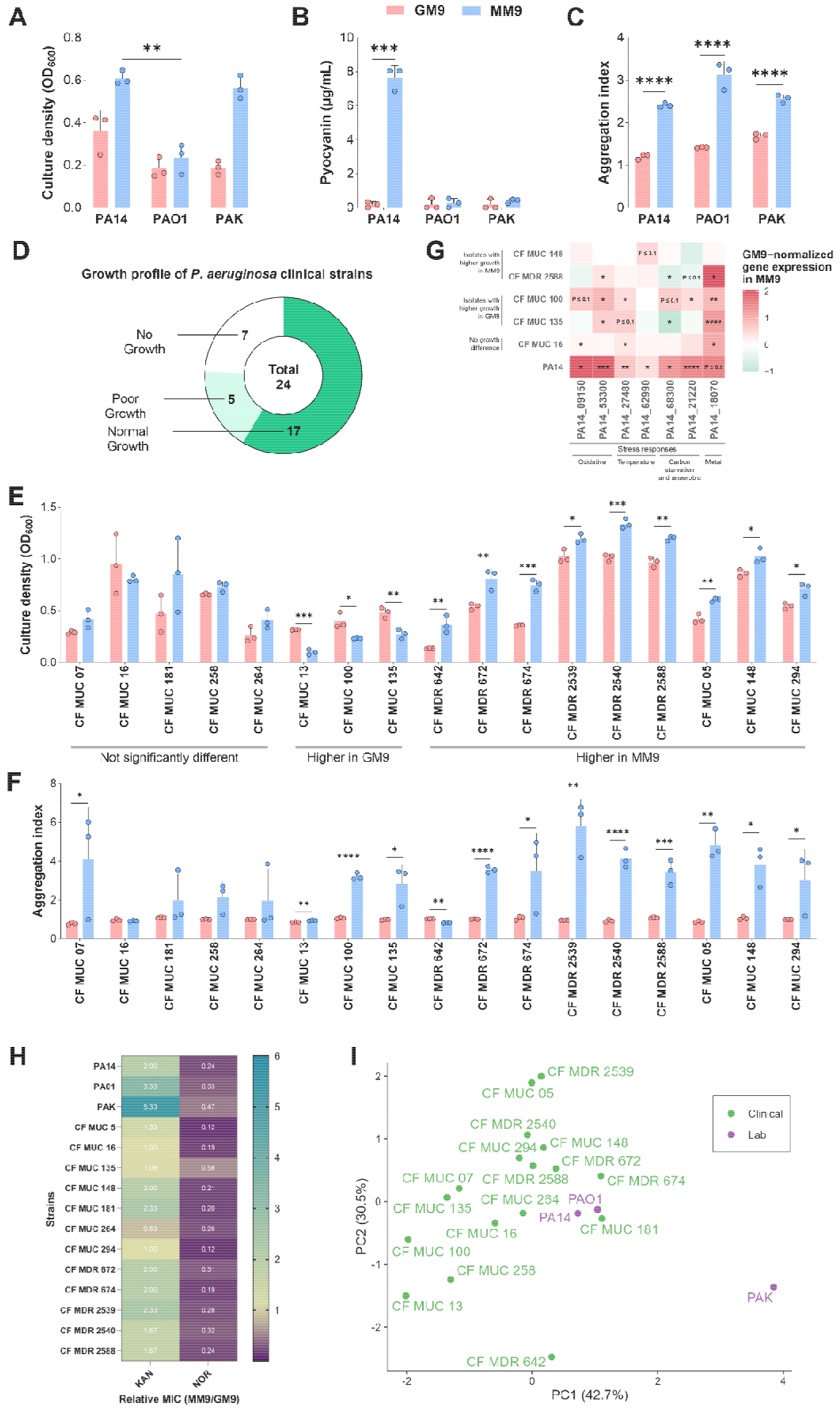
Phenotypes of *P. aeruginosa* lab and clinical strains grown in MM9 and GM9. (A-C) Lab strains growth phenotypes. **(A)** OD_600_ was measured for all the three lab strains after 16 hours of incubation at 37°C. **(B)** Pyocyanin was extracted from the supernatant, then quantified as described in the methods. **(C)** Aggregation index was calculated as the ratio of OD_600_ post-sonication to OD_600_ pre-sonication of 24-old bacterial culture. **(D-F)** Clinical cystic fibrosis (CF) isolates growth phenotypes. **(D)** Proportion of clinical strains that robustly grew in GM9 and MM9 (Normal growth), poorly grew (OD_600_ below average, 0.5), or did not grew (No growth). **(E)** Culture density based on OD_600_ measurement after 16 hours of incubation at 37°C. **(F)** Aggregation index was calculated as the ratio of OD_600_ post-sonication to OD_600_ pre-sonication of 48- old bacterial culture. **(G)** Gene expression of selected stress responsive genes in clinical strains. PA14 and the clinical strains were grown overnight in GM9 and MM9 before their RNA was extracted, followed by qRT-PCR. Expression values were normalized to a reference housekeeping gene, i.e., the 16S ribosomal RNA gene *PA14_08570*. Shown expression values in MM9 were normalized to that of GM9. Color indicates the average of biological triplicates. **(H)** Minimum inhibitory concentration (MIC) of KAN and NOR against *P. aeruginosa* strains in MM9 vs. GM9. The MIC values for KAN and NOR in MM9 and GM9 were determined through a broth microdilution assay, and the MIC values for each antibiotic in MM9 were normalized to the corresponding antibiotic in the GM9 medium. CF: cystic fibrosis, MUC: mucoid strains, MDR: multidrug-resistant strain, KAN: Kanamycin, NOR: Norfloxacin. Values represent the means of three independent experiments ± standard deviation. Statistical significance was calculated by two-tailed unpaired t-test. * P < 0.05, ** P < 0.01, *** P < 0.001, and **** P < 0.0001.

**Table 5.** Description of lab strains used in this study.

| Strain | Description | Reference |
| --- | --- | --- |
| PA14 | *P. aeruginosa* strain UCBPP-PA14; prototrophic; clinical isolate from burn patient | [83] |
| PAK | *P. aeruginosa* strain K; hyperpiliated; sensitivity to Pf1 phage; prototrophic | [62, 84] |
| PAO1 | *P. aeruginosa* strain; prototrophic; clinical isolate from a wound | [61] |
| JE2 | *S. aureus* MRSA strain USA300 LAC cured of plasmids | [85] |

Encouraged by our previous results and the knowledge that malonate decarboxylase protein is abundant in a clinical isolate of *P. aeruginosa* [64], we wondered if the previously described phenotypes can be observed in other clinical strains of *P. aeruginosa*, isolated from lungs of cystic fibrosis (CF) patients. To examine this, we utilized a collection of 24 clinical isolates of *P. aeruginosa*, which included strains showing multidrug resistance or mucoid phenotypes **(Table 6)**. First, we examined the growth phenotype for all clinical isolates in MM9 and GM9. While the majority (17 isolates) were able to grow in GM9 and MM9, five isolates grew poorly, and two showed no growth **(Figure 4D).** More than half of the clinical isolates showed more robust growth in MM9 rather than in GM9, similar to what we previously observed for PA14 **(Figure 4E)**. Only three isolates showed higher growth in GM9 vs. MM9 **(Figure 4E)**. Next, we qualitatively assessed the blue-green pigmentation of the clinical strains grown in MM9, as an indication of the induced pyocyanin production **(Table 6)**. Most of the clinical strains showing blue-green pigmentation had better growth in MM9. We were finally interested in examining the aggregation phenotype among our clinical isolate collection. We found that aggregation is a common outcome of malonate- utilization among the isolates (around two-thirds showed high aggregation index) **(Figure 4F)**.

**Table 6.** Clinical strains used in this study and their phenotypes. Blue-green pigmentation was qualitatively assessed by visually examining bacterial cultures.

| Strain | Clinical phenotype | Blue-green pigment |
| --- | --- | --- |
| CF MUC 05 | Mucoid | - |
| CF MUC 07 | Mucoid | - |
| CF MUC 13 | Mucoid | - |
| CF MUC 16 | Mucoid | - |
| CF MUC 19 | Mucoid | - |
| CF MUC 20 | Mucoid | - |
| CF MUC 100 | Mucoid | - |
| CF MUC 135 | Mucoid | - |
| CF MUC 148 | Mucoid | + |
| CF MUC 181 | Mucoid | + |
| CF MUC 221 | Mucoid | - |
| CF MUC 258 | Mucoid | - |
| CF MUC 264 | Mucoid | - |
| CF MUC 294 | Mucoid | - |
| CF MDR 490 | Multidrug resistant | - |
| CF MDR 642 | Multidrug resistant | - |
| CF MDR 666 | Multidrug resistant | - |
| CF MDR 672 | Multidrug resistant | + |
| CF MDR 674 | Multidrug resistant | + |
| CF MDR 2350 | Multidrug resistant | - |
| CF MDR 2539 | Multidrug resistant | + |
| CF MDR 2540 | Multidrug resistant | + |
| CF MDR 2577 | Multidrug resistant | - |
| CF MDR 2588 | Multidrug resistant | + |

After observing that many of the clinical isolates show similar phenotypes to that of PA14, we wondered if we can also observe similar trends at the transcriptional level. To evaluate this, we selected five clinical strains that showed variable growth phenotypes in MM9 and GM9 and examined their expression of stress response genes. Briefly, MUC 16 was selected as it showed no significant growth difference in MM9 and GM9, MDR 2588 and MUC 148 showed higher growth in MM9, and MUC 135 and MUC 100 showed higher growth in GM9. We examined the expression of genes related to stress response that we identified to be upregulated in MM9. These included genes belonging to metal, oxidative, anaerobic, and temperature stress responses. While we observed a significant variability in the gene expression of the selected genes, a few showed similar trends **(Figure 4G)**. Specifically, genes involved in metal and oxidative stress response (*PA14_53300* and *PA14_18070*) were consistently upregulated in MM9 vs. GM9 in four out of the five selected clinical isolates, regardless of their growth preference **(Figure 4G)**, similar to what was previously observed in PA14. In general, malonate utilization appears to be linked to comparable stress response gene expression patterns and phenotypes among numerous clinical isolates and laboratory strains of *P. aeruginosa*.

Since one of our notable previous observations was that PA14 displayed differential susceptibility to antibiotics we next wanted to test if this is also conserved across all tested strains. Specifically, we wanted to examine if both clinical and lab strains exhibit increased tolerance to kanamycin and greater susceptibility to norfloxacin in MM9 compared to GM9 [65, 66]. We examined the impact of malonate and glycerol as carbon sources on antibiotic susceptibility using a broth microdilution assay for both kanamycin (up to 1200 μg/ml) and norfloxacin (up to 100 μg/ml).

In line with the lab strains, most clinical isolates (CF MUC 5, CF MUC 148, CF MUC 181, CF MUC 264, CF MDR 672, CF MDR 674, CF MDR 2539, CF MDR 2540, and CF MDR 2588) exhibited increased tolerance to kanamycin in MM9 compared to GM9 **(Figure 4H and Table 7)**. However, unlike control strains, CF MUC 13 and CF MUC 100 displayed heightened kanamycin tolerance in GM9 compared to MM9. Furthermore, several tested strains maintained consistent susceptibility or tolerance to kanamycin regardless of the carbon source. CF MUC 7 did not grow in MM9 but thrived in GM9. Interestingly, CF MUC 7 did not grow in GM9 when glacial acetic acid was used as a vehicle control. In addition, all strains demonstrated greater tolerance to norfloxacin in GM9 media than MM9 **(Figure 4H and Table 7)**. In summary, our results indicate that both lab and clinical *P. aeruginosa* strains tend to exhibit increased tolerance to kanamycin and heightened susceptibility to norfloxacin when malonate is utilized as a sole carbon source.

**Table 7.** MIC of Kanamycin (KAN) and norfloxacin (NOR) against wild-type and clinical isolates of *P. aeruginosa* in MM9 and GM9 media. ng: no growth, ng-aa: no growth with glacial acetic acid as vehicle control.

| Strain | MIC (µg/ml) |  |  |  |
| --- | --- | --- | --- | --- |
|  | Kanamycin |  | Norfloxacin |  |
|  | GM9 | MM9 | GM9 | MM9 |
| PA14 | 150-300 | 300-600 | 0.78-0.39 | 0.09-0.19 |
| PAO1 | 75-150 | 300 | 6.25-12.5 | 1.56-3.12 |
| PAK | 150-300 | 600-1200 | 0.19 | 0.09 |
| CF MUC 5 | 600 | 600-1200 | 1.56-3.12 | 0.09-0.78 |
| CF MUC 7 | 1200 | ng | ng-aa | ng |
| CF MUC 13 | 1200->1200 | 600->1200 | ng-aa-0.78 | ng-0.09 |
| CF MUC 16 | >1200 | >1200 | 25-100 | 6.25 |
| CF MUC 100 | >1200 | 600->1200 | 0.39->100 | 0.19->100 |
| CF MUC 135 | 300->1200 | 300->1200 | 0.39-6.25 | 0.09-3.12 |
| CF MUC 148 | 300-600 | 600-1200 | 1.56 | 0.19-0.39 |
| CF MUC 181 | 300-600 | 600-1200 | 1.56 | 0.19-0.39 |
| CF MUC 264 | >1200 | 600->1200 | 6.25-12.5 | 0.39-6.25 |
| CF MUC 294 | >1200 | >1200 | 12.5 | 1.56 |
| CF MDR 672 | 300 | 600 | 0.19-0.39 | 0.09 |
| CF MDR 674 | 600 | 1200 | 0.39-0.78 | 0.09 |
| CF MDR 2539 | 300-600 | 600-1200 | 1.56 | 0.39-0.78 |
| CF MDR 2540 | 600 | 600-1200 | 0.19-0.39 | 0.09-0.39 |
| CF MUC 258 | ng | ng | ng | ng |
| CF MDR 2588 | 300 | 300-600 | 0.39-0.78 | 0.09-0.19 |

Finally, based on the phenotypes we have examined so far across all strains, i.e., growth density, aggregation, and tolerance to antibiotics, we were curious to see whether the phenotypes for the tested lab strains would have an overall closer resemblance with each other versus the tested clinicals strains. To this end, we compiled the aforementioned datasets and performed a principal component analysis (PCA). Interestingly, two of the lab strains (i.e., PA14 and PAO1) clustered closely to each other and to other clinicals strains, unlike the lab strain PAK **(Figure 4I).** The more distant phenotype of PAK is due to the higher growth density ratio (MM9/GM9) of PAK **(Figure 4A)** and tolerance ratio (MM9/GM9) to kanamycin **(Figure 4H)** in MM9 in comparison with other strains. Overall, this suggests that several of the malonate- utilization-associated phenotypes are conserved across both lab and clinical strains.

## Discussion

Bacteria occupying different niches can utilize different carbon sources. This can in turn have a profound impact on their growth rate, virulence, phenotypes, and resistance to antibiotics [9, 67–70]. Earlier studies have described the role of carbon sources in the metabolism of *P. aeruginosa* [71–74]. Our study highlights the role of malonate as a sole carbon source in inducing a global stress response in *P. aeruginosa*. It is known that the differential activation of the glyoxylate cycle in the presence of different carbon sources can affect both the pathogenesis and virulence of *P. aeruginosa* [75]. Our transcriptomic data pointed out that, in the presence of malonate, *P. aeruginosa* implements specific pathways to survive and adapt to its environment, modifying its virulence and stress response. In our study, we found that both the glyoxylate and methyl citrate cycle-associated genes were upregulated in the presence of malonate. The transcriptomic data further highlighted the intimate interaction between malonate utilization and various stress responsive genes in *P. aeruginosa*. Our transcriptomic data showed an upregulation in genes associated with metal stress response. This intrigued us to examine the metal ions accumulation in the presence of malonate. Aluminum and strontium were the two main metals that showed a significant accumulation in MM9 vs. GM9. We speculate that aluminum accumulation could be the result of differential siderophore production [76]. While we expected to observe copper accumulation in MM9, we observed none. We speculate that as an adaptation strategy, *P. aeruginosa* eliminates excess copper through the production and secretion of the copper-containing small molecule fluopsin C [57]. This is supported by the remarkable upregulation of the fluopsin C-biosynthetic genes.

Because *P. aeruginosa* and *S. aureus* are known to cause chronic infections, including lung infections, we were curious if malonate influences their interspecies interaction. A recent study demonstrated that the ability of *P. aeruginosa* to break down acetoin induces trophic cooperation between *S. aureus* and *P. aeruginosa* and improves their survival [77]. Our experiments revealed that malonate-utilization allowed *P. aeruginosa* to outcompete *S. aureus*. This could be explained by our previous observations showing that malonate induces the expression of biosynthetic genes of fluopsin C and pyocyanin, both of which are toxic to *S. aureus* [57, 58, 78]. We excluded the possibility that acetoin could be involved here because our transcriptomic data showed an upregulation in metabolic genes of acetoin, which should be improving *S. aureus* fitness, but was not observed here. However, other virulence factors not examined in our study could be involved here. This suggests that in a niche where malonate is available as a carbon source, it can allow *P. aeruginosa* to outcompete *S. aureus*.

Our results allowed us to postulate if phenotypes associated with malonate- utilization were similar in other lab and clinical strains of *P. aeruginosa*. Interestingly, most of the tested strains demonstrated better growth in MM9 vs. GM9. This result is of high clinical relevance because traditional lab medium for selective growth of *P. aeruginosa* is *Pseudomonas* Isolation Agar (PIA), which requires glycerol to be added to it (2%) to serve as an energy source and to enhance pyocyanin production (BD Diagnostics or Remel). If strains of *P. aeruginosa* do not utilize glycerol as a carbon source, this can hinder isolating such strains that may utilize another carbon source such as malonate or mischaracterize *P. aeruginosa* as malonate is a much better enhancer of pyocyanin production than glycerol [14]. This is further supported by previous studies that investigated nutrient availability in sputum samples. For example, a study showed that malonate is among the abundant metabolites in the sputum of healthy volunteers [17]. However, it is not known if malonate is more or less abundant in the sputum of cystic fibrosis patients, which is an important issue that warrants further investigation. Therefore, our results highlight the importance of studying the nutrient adaptation of different strains of *P. aeruginosa* to assist in their identification and isolation. Furthermore, we showed that the expression of certain genes involved in stress response are similarly overexpressed in clinical strains of *P. aeruginosa.* Overall, our findings show that clinical strains have unique transcriptional features that may deeply impact *P. aeruginosa’*s adaptation to specific environments found inside the host. Further genomic investigations are required on more clinical isolates to decipher potential mechanisms that facilitate their adaptation in response to different carbon sources. Further research on the metabolomics and proteomics analysis of *P. aeruginosa* exposed to malonate will provide us with valuable insights into the specific metabolites and proteins that are generated or triggered in the presence of this particular carbon source. Such pathways can eventually be used as a potential therapeutic target to combat *Pseudomonas*-associated infections and help us identify new strategies complementary or alternatives to antibiotics to efficaciously combat this notorious pathogen.

Our present findings align with the results obtained in our previous work, demonstrating increased and reduced tolerance to kanamycin and norfloxacin, respectively, in MM9 compared to GM9 [65, 66]. In this latest investigation, we increased the spectrum of our lab strains and incorporated clinical strains to reinforce and validate our earlier observations. Most clinical strains, including the three lab control strains, exhibited heightened tolerance to kanamycin in MM9 compared to GM9.

Conversely, all clinical strains and the three lab strains, demonstrated increased susceptibility to norfloxacin in MM9 as opposed to GM9.

Overall, we present malonate as an important carbon source to *P. aeruginosa* that can influence its carbon metabolism, elicit global stress response, and lead to intracellular metal accumulation, improving its pathogenicity against other bacterial competitors. Furthermore, phenotypes associated with malonate-utilization are not specific to lab strains of *P. aeruginosa*, but also to clinical strains that warrant further investigation.

## Material and Methods

### Bacterial growth and strains

Bacterial strains used in this study are described in Tables 2 and 3. We used the strain PA14 and its isogenic *mariner* transposon mutants (**Supplementary Table 1**) for the majority of the experiments. Bacterial strains were routinely grown overnight in Luria-Bertani (LB) broth. When needed, antibiotics were added at the following concentrations: 50 μg/mL of ampicillin and 15 μg/mL of gentamicin.

For analysis of the effect of malonate as a sole carbon source on the growth and virulence of PA14, we used the established M9 minimal medium (6.0 g Na_2_HPO_4_, 3.0 g KH_2_PO_4_, 0.5 g NaCl, 1.0 g NH_4_Cl per L supplemented with 0.2 mM CaCl_2_ and 2 mM MgSO_4_) (Fisher Scientific) as a basal medium [79]. No iron source was added to the M9 minimal medium. For the control medium, we modified the M9 by adding 1% glycerol (v/v; 110 mM) as a sole carbon source (GM9). We previously utilized this concentration of glycerol as a carbon source in analyzing the regulation of *P. aeruginosa* virulence genes [14]. For M9 media containing malonate as a sole carbon source (MM9), we added 40 mM malonate (or 100 mM malonate if explicitly mentioned), as sodium malonate dibasic (MilliporeSigma, St. Louis, MO). For analysis of gene expression and virulence factor production, PA14 was routinely grown in GM9 and MM9 for 16 hours.

### RNA extraction

PA14 in MM9 and GM9 media was grown overnight for 16 hours at 37°C in shaking condition. The overnight grown culture was then centrifuged at 5,000 rpm speed for 10 min. After discarding the supernatant, bacterial pellets were lysed by the addition of lysozyme and proteinase K for 15 min at room temperature. RNA was extracted using the RNeasy Mini Kit (Qiagen) according to the manufacturer’s protocol. RNA solution was digested with the RNase-free DNase set (Qiagen), followed by on- column DNase digestion to eliminate any remaining traces of genomic DNA. The purified RNA was quantified using a NanoDrop spectrophotometer (NanoDrop Technologies, Wilmington, DE). The samples were then sent to Genewiz for library prep and Illumina HiSeq. Only samples with an RNA integration number greater than 8.0 were used for cDNA library preparation.

### Transcriptomic (RNA-seq) analysis

RNA-seq data were analyzed using Rockhopper software implementing reference-based transcript assembly with UCBPP-PA14 as a reference genome followed by calculating the fold change for the transcripts at each growth condition [18]. NCBI Reference Sequence NC_008463.1 was used. Datasets were normalized using upper quartile normalization, then transcript abundance was quantified using reads assigned per the kilobase of target per million mapped reads normalization method. The selection criteria for differential expression required genes to have a fold change of ≥ 1.5 and a Q value of ≤ 0.05 to be considered significant. The Q value was obtained by adjusting the P value using the Benjamini–Hochberg procedure.

### Real-Time Quantitative Reverse Transcription PCR (qRT-PCR)

One μL of RNA was used for cDNA synthesis. For qRT-PCR, equal amounts of cDNA were mixed with iQ SYBR Green Supermix (Bio-Rad, Hercules, CA) together with 2 μM of specific primers for each gene examined. Amplification and detection were done using CFX96 Deep Well Real-Time PCR System (Bio-Rad) and analysis of gene expression was done using CFX Manager 3.1 software (Bio-Rad). Each experiment consisted of three biological replicates analyzed in triplicate. Quantity of cDNA in the samples was normalized using the 16S ribosomal RNA gene *PA14_08570*. List of primers used in this study are mentioned in **Supplementary Table 2**.

### Pyocyanin measurement

This was performed following previously described protocols [63, 80]. 2 mL samples of the supernatant fractions of cultures of PA14, PAO1 and PAK grown in GM9 or MM9 for 16 hours were isolated and mixed with chloroform (1:1). The lower layer was then separated and mixed with two ml of 0.2 N HCl. The pyocyanin-rich organic upper layer which is pink in color was then extracted, and the absorbance at 520 nm was determined. Values were normalized by dividing by the OD_600_ of each culture prior application of the following formula. The amount of pyocyanin, in μg/mL, was calculated using the following formula: OD_520_ × 17.072 = μg of pyocyanin/mL [63].

### Calculation of the aggregation index

The aggregation index was calculated as the ratio of post-sonication OD_600_ to pre-sonication OD_600_ of overnight cultures [81].

### Intracellular metal ions measurement using ICP-MS

Samples were prepared for ICP-MS following an established protocol with slight modifications [54]. Bacterial cells were cultured overnight in Chelex-treated MM9 and GM9 for 16 hours in the presence of a cocktail of essential metal ions add to the media (Mg 1mM, Mn 25 µM, Ca 100 µM, Fe 5 µM, Zn 25 µM, Co 0.1 µM, and Cu 0.1 µM).

These cultures were pelleted and washed in Chelex-treated 500 μL phosphate-buffered saline (PBS). Next, the pellet was suspended in 1 mL metal-free water (Chelex treated) and sonicated at 100% amplitude for 30 seconds each. The probe was washed with Chelex-treated water and the control solution used was the sonicated water. The probe was wiped and rinsed between samples as thoroughly as possible using water fresh from our filtration system. We sonicated 1 water sample between every 2 samples to account for metal fluctuations that may occur over the sonication process. Samples were normalized to equal protein concentration and 100 μL of each sample was digested by adding 1 mL 50% HNO3 (Optima grade; Fisher) followed by overnight incubation at 50°C in a metal-free 15-ml conical tube. After digestion, samples were diluted to a 20-ml final volume in Milli-Q water and submitted for inductively coupled plasma mass spectrometry (ICP-MS) analysis. Levels of Ag, Al, As, B, Be, Cd, Co, Cr, Cs, Cu, Fe, Ga, Hg, K, Li, Mg, Mn, Na, Ni, Pb, Rb, Se, Sr, Ti, U, V and Zn were measured. Quality analysis and quality control included Rh-103 and In-115 as internal standards as well as blanks and continuing calibration checks which were run every 10 samples. Calibration standards were prepared in 50 mL polypropylene tubes with 2% HNO_3_ to match the sample matrix. Detection limits for the elements quantified were: Li = Cu = 0.01 ppb, Be = 0.0006 ppb, Mg = Al = 0.47 ppb, V = 0.04 ppb, Cr = Mn = Cs = Rb = U = 0.001 ppb, Fe = 0.04 ppb, Co = 0.005 ppb, Ni = Sr = 0.02 ppb, Ga = 0.002, Zn = 0.11 ppb). Ion levels were normalized to the endogenous ion levels of the control samples.

### Co-culture in MM9 and GM9 media

Prior to inoculation of cultures into MM9 and GM9 growth medium, the laboratory reference strain of *P. aeruginosa* UCBPP-PA14 was cultured in LB and the wild-type strain of *S. aureus*, USA300 JE2 was cultured in Tryptic soy broth (TSB). Overnight cultures were washed thrice with filter-sterilized PBS to remove media contamination from either LB or TSB. After a final wash with M9, cells were normalized to an OD_600_ of 1.0 (culture density of ∼10^8^). Normalized cells were then suspended in MM9 and GM9 as a mono-culture or co-culture. Cells were then incubated as monocultures or co-cultures for 24 hours at 37°C under shaking condition. Following incubation, bacterial cells were diluted in sterile, PBS and plated on selective media; *P. aeruginosa* monocultures were plated on Pseudomonas Isolation Agar (PIA) plates, *S. aureus* monocultures were plated on Mannitol Salt Agar (MSA) plates, and co-cultures were plated on both plates to observe differences in microbial growth.

### Antibiotic susceptibility testing for bacterial strains

Minimum inhibitory concentration (MICs) of kanamycin and norfloxacin against wild-type and clinical isolates of *P. aeruginosa* was determined in both MM9 and GM9 using broth microdilution method [82]. Overnight cultures were adjusted to an optical density at 600 nm of 1.0 in 1X M9 and diluted to ∼5X10^5^ colony-forming units (CFU) per ml. Both kanamycin and norfloxacin were diluted in 1X M9, then added to 96-well plates containing bacterial inoculum in either MM9 or GM9. Cells without any drug were also incubated as a positive growth control. Distilled water and glacial acetic acid were added as vehicle controls for kanamycin and norfloxacin, respectively, as the initial drug stocks were prepared. Plates were incubated at 37°C for 24 hours at static condition. Bacterial growth was assessed using 20µL of 0.2 mg/mL resazurin solutions in each well. Following another 12 hours of incubation with resazurin, the MIC was determined visually by observing the transition from blue (non-viable) to pink (viable) color. We standardized the MIC data in MM9 against GM9 and represented the results in a heatmap for comparative analysis. A value of 1, denoted by a yellow color in the heatmap, indicated that a strain remained unaffected by the action of any tested antibiotic in both media. Increasing tolerance to a specific antibiotic in MM9 was represented by shades of light to deep green, while greater susceptibility was depicted by shades of light to deep purple.

### Principal component analysis (PCA) of quantitative phenotypes across strains

The following phenotypic datasets, i.e., culture growth density, aggregation index, and MIC of kanamycin and norfloxacin, in MM9 normalized to those of GM9. The average of replicates per phenotypic dataset per strain was calculated. Missing values were imputed to 1. Each dataset was scaled and centered before principal component analysis.

### Statistical Analysis

Statistical analyses and graphics plotting were performed using R 4.0.5 and GraphPad Prism 9.0 (GraphPad Software, Inc., San Diego, CA). Unpaired t-test was used to determine statistical significance (unless otherwise stated).

### Data availability

The raw sequencing data of each sample have been deposited under BioProject accession number PRJNA862157 in the National Center for Biotechnology Information (NCBI) BioProject database.

## Acknowledgements

This work was supported by NIH/NIGMS grant R15GM128072 and NIH/NIAID grant R01AI173686 to CAW and by the Burn Center of Research Excellence (BCoRE) in the Department of Surgery at TTUHSC, Lubbock, TX to ANH and MME. HAM, MME, and KB were supported by the Doctoral Dissertation Completion Fellowships granted from Texas Tech University Graduate School, Lubbock, TX.

## References

1. Van Delden, C. and B.H. Iglewski, Cell-to-cell signaling and Pseudomonas aeruginosa infections. Emerging infectious diseases, 1998. 4(4): p. 551–560.

2. Driscoll, J.A., S.L. Brody, and M.H. Kollef, The Epidemiology, Pathogenesis and Treatment of Pseudomonas aeruginosa Infections. Drugs, 2007. 67(3): p. 351–368.

3. Arai, H., Regulation and Function of Versatile Aerobic and Anaerobic Respiratory Metabolism in Pseudomonas aeruginosa. Frontiers in Microbiology, 2011. 2: p. 103.

4. Cases, I. and V. de Lorenzo, Promoters in the environment: transcriptional regulation in its natural context. Nature Reviews Microbiology, 2005. 3(2): p. 105–118.

5. Meylan, S., et al., Carbon Sources Tune Antibiotic Susceptibility in Pseudomonas aeruginosa via Tricarboxylic Acid Cycle Control. Cell chemical biology, 2017. 24(2): p. 195–206.

6. Elmassry, M.M., et al., Pseudomonas aeruginosa Alters Its Transcriptome Related to Carbon Metabolism and Virulence as a Possible Survival Strategy in Blood from Trauma Patients. mSystems, 2019. 4(4): p. e00312–18.

7. Bartell, J.A., et al., Reconstruction of the metabolic network of Pseudomonas aeruginosa to interrogate virulence factor synthesis. Nature Communications, 2017. 8(1): p. 14631.

8. Rojo, F., Carbon catabolite repression in Pseudomonas: optimizing metabolic versatility and interactions with the environment. FEMS Microbiology Reviews, 2010. 34(5): p. 658–684.

9. Troitzsch, A., et al., Carbon Source-Dependent Reprogramming of Anaerobic Metabolism in Staphylococcus aureus. Journal of Bacteriology, 2021. 203(8): p. e00639–20.

10. Maderbocus, R., et al., Crystal structure of a Pseudomonas malonate decarboxylase holoenzyme hetero-tetramer. Nature Communications, 2017. 8(1): p. 160.

11. Dimroth, P. and H. Hilbi, Enzymic and genetic basis for bacterial growth on malonate. Molecular Microbiology, 1997. 25(1): p. 3–10.

12. Wang, Z., et al., A Disjointed Pathway for Malonate Degradation by Rhodopseudomonas palustris. Applied and Environmental Microbiology, 2020. 86(11): p. e00631–20.

13. Karunakaran, R., K. East Alison, and S. Poole Philip, Malonate Catabolism Does Not Drive N2 Fixation in Legume Nodules. Applied and Environmental Microbiology, 2013. 79(14): p. 4496–4498.

14. Elmassry, M.M., et al., Malonate utilization by Pseudomonas aeruginosa affects quorum- sensing and virulence and leads to formation of mineralized biofilm-like structures. Mol Microbiol, 2021. 116(2): p. 516–537.

15. Li, X., et al., Spatial patterns of carbonate biomineralization in biofilms. Appl Environ Microbiol, 2015. 81(21): p. 7403–10.

16. Reichhardt, C. and M.R. Parsek, Confocal Laser Scanning Microscopy for Analysis of Pseudomonas aeruginosa Biofilm Architecture and Matrix Localization. Front Microbiol, 2019. 10: p. 677.

17. Farne, H., et al., Comparative Metabolomic Sampling of Upper and Lower Airways by Four Different Methods to Identify Biochemicals That May Support Bacterial Growth. Front Cell Infect Microbiol, 2018. 8: p. 432.

18. Tjaden, B., A computational system for identifying operons based on RNA-seq data. Methods, 2020. 176: p. 62–70.

19. Thaden, J.T., S. Lory, and T.S. Gardner, Quorum-sensing regulation of a copper toxicity system in Pseudomonas aeruginosa. Journal of bacteriology, 2010. 192(10): p. 2557–2568.

20. Teitzel, G.M., et al., Survival and Growth in the Presence of Elevated Copper: Transcriptional Profiling of Copper-Stressed Pseudomonas aeruginosa. Journal of Bacteriology, 2006. 188(20): p. 7242.

21. Imlay, J.A., The molecular mechanisms and physiological consequences of oxidative stress: lessons from a model bacterium. Nature Reviews Microbiology, 2013. 11(7): p. 443–454.

22. Imlay, J.A., The mismetallation of enzymes during oxidative stress. The Journal of biological chemistry, 2014. 289(41): p. 28121–28128.

23. Lu, S., et al., CDD/SPARCLE: the conserved domain database in 2020. Nucleic acids research, 2020. 48(D1): p. D265–D268.

24. Maura, D., et al., Evidence for Direct Control of Virulence and Defense Gene Circuits by the Pseudomonas aeruginosa Quorum Sensing Regulator, MvfR. Scientific Reports, 2016. 6(1): p. 34083.

25. Palma, M., et al., Transcriptome Analysis of the Response of Pseudomonas aeruginosa to Hydrogen Peroxide. Journal of Bacteriology, 2004. 186(1): p. 248.

26. Aharoni, N., et al., Gene expression in Pseudomonas aeruginosa exposed to hydroxyl- radicals. Chemosphere, 2018. 199: p. 243–250.

27. Hunter, R.C. and D.K. Newman, *A putative ABC transporter,* hatABCDE, is among molecular determinants of pyomelanin production in Pseudomonas aeruginosa. J Bacteriol, 2010. 192(22): p. 5962–71.

28. Hunter, R.C. and D.K. Newman, *A putative ABC transporter,* hatABCDE, is among molecular determinants of pyomelanin production in Pseudomonas aeruginosa. Journal of bacteriology, 2010. 192(22): p. 5962–5971.

29. Kojic, M., C. Aguilar, and V. Venturi, TetR Family Member PsrA Directly Binds the Pseudomonas rpoS and psrA Promoters. Journal of Bacteriology, 2002. 184(8): p. 2324.

30. Basta, D.W., M. Bergkessel, and D.K. Newman, Identification of Fitness Determinants during Energy-Limited Growth Arrest in Pseudomonas aeruginosa. mBio, 2017. 8(6): p. e01170–17.

31. Liang, P., et al., The aerobic respiratory chain of Pseudomonas aeruginosa cultured in artificial urine media: Role of NQR and terminal oxidases. PloS one, 2020. 15(4): p. e0231965–e0231965.

32. Filiatrault, M.J., et al., Identification of Pseudomonas aeruginosa genes involved in virulence and anaerobic growth. Infection and immunity, 2006. 74(7): p. 4237–4245.

33. Torres, A., et al., NADH Dehydrogenases in Pseudomonas aeruginosa Growth and Virulence. Frontiers in Microbiology, 2019. 10: p. 75.

34. McPhee, J.B., et al., The major outer membrane protein OprG of Pseudomonas aeruginosa contributes to cytotoxicity and forms an anaerobically regulated, cation-selective channel. FEMS Microbiology Letters, 2009. 296(2): p. 241–247.

35. Boes, N., K. Schreiber, and M. Schobert, SpoT-Triggered Stringent Response Controls Gene Expression in Pseudomonas aeruginosa. Journal of Bacteriology, 2008. 190(21): p. 7189.

36. Sønderholm, M., et al., Pseudomonas aeruginosa Aggregate Formation in an Alginate Bead Model System Exhibits In Vivo-Like Characteristics. Applied and environmental microbiology, 2017. 83(9): p. e00113-17.

37. Palmer, K.L., S.A. Brown, and M. Whiteley, Membrane-bound nitrate reductase is required for anaerobic growth in cystic fibrosis sputum. Journal of bacteriology, 2007. 189(12): p. 4449–4455.

38. Comolli, J.C. and T.J. Donohue, Differences in two Pseudomonas aeruginosa cbb3 cytochrome oxidases. Molecular Microbiology, 2004. 51(4): p. 1193–1203.

39. Clay, M.E., et al., Pseudomonas aeruginosa lasR mutant fitness in microoxia is supported by an Anr-regulated oxygen-binding hemerythrin. Proceedings of the National Academy of Sciences of the United States of America, 2020. 117(6): p. 3167–3173.

40. Arai, H., M. Mizutani, and Y. Igarashi, *Transcriptional regulation of the nos genes for nitrous oxide reductase in Pseudomonas aeruginosa.* Microbiology (Reading, England), 2003. 149(Pt 1): p. 29–36.

41. Trunk, K., et al., Anaerobic adaptation in Pseudomonas aeruginosa: definition of the Anr and Dnr regulons. Environmental Microbiology, 2010. 12(6): p. 1719–1733.

42. Arat, S., G.S. Bullerjahn, and R. Laubenbacher, A Network Biology Approach to Denitrification in Pseudomonas aeruginosa. PLOS ONE, 2015. 10(2): p. e0118235.

43. Zhu, X., et al., Nitric Oxide-Mediated Induction of Dispersal in Pseudomonas aeruginosa Biofilms Is Inhibited by Flavohemoglobin Production and Is Enhanced by Imidazole. Antimicrobial Agents and Chemotherapy, 2018. 62(3): p. e01832–17.

44. Zimmermann, A., et al., Anaerobic growth and cyanide synthesis of Pseudomonas aeruginosa depend on anr, a regulatory gene homologous with fnr of Escherichia coli. Molecular Microbiology, 1991. 5(6): p. 1483–1490.

45. Cao, P., et al., A Pseudomonas aeruginosa small RNA regulates chronic and acute infection. Nature, 2023. 618(7964): p. 358-364.

46. Schiessl, K.T., et al., Phenazine production promotes antibiotic tolerance and metabolic heterogeneity in Pseudomonas aeruginosa biofilms. Nature Communications, 2019. 10(1): p. 762.

47. Sporer, A.J., et al., *Pseudomonas aeruginosa PumA acts on an endogenous phenazine to promote self-resistance.* Microbiology (Reading, England), 2018. 164(5): p. 790–800.

48. Smith, P., et al., Bacterial Cheaters Evade Punishment by Cyanide. iScience, 2019. 19: p. 101-109.

49. Yan, H., et al., Conditional quorum-sensing induction of a cyanide-insensitive terminal oxidase stabilizes cooperating populations of Pseudomonas aeruginosa. Nature Communications, 2019. 10(1): p. 4999.

50. Koizumi, S., K. Suzuki, and S. Yamaguchi, Heavy metal response of the heat shock protein 70 gene is mediated by duplicated heat shock elements and heat shock factor 1. Gene, 2013. 522(2): p. 184–191.

51. Tran, T.D.-H., et al., Decrease in penicillin susceptibility due to heat shock protein ClpL in Streptococcus pneumoniae. Antimicrobial agents and chemotherapy, 2011. 55(6): p. 2714–2728.

52. Basta, D.W., et al., Heat-shock proteases promote survival of Pseudomonas aeruginosa during growth arrest. Proc Natl Acad Sci U S A, 2020. 117(8): p. 4358–4367.

53. Hinz, A., et al., Membrane proteases and aminoglycoside antibiotic resistance. Journal of bacteriology, 2011. 193(18): p. 4790–4797.

54. Wakeman, C.A., et al., Differential activation of Staphylococcus aureus heme detoxification machinery by heme analogues. Journal of Bacteriology, 2014. 196: p. 1335–1342.

55. Schalk, I.J., M. Hannauer, and A. Braud, New roles for bacterial siderophores in metal transport and tolerance. Environmental Microbiology, 2011. 13(11): p. 2844–2854.

56. Braud, A., et al., Presence of the siderophores pyoverdine and pyochelin in the extracellular medium reduces toxic metal accumulation in Pseudomonas aeruginosa and increases bacterial metal tolerance. Environmental Microbiology Reports, 2010. 2(3): p. 419–425.

57. Patteson, J.B., et al., Biosynthesis of fluopsin C, a copper-containing antibiotic from Pseudomonas aeruginosa. Science, 2021. 374(6570): p. 1005-1009.

58. Hall, S., et al., Cellular Effects of Pyocyanin, a Secreted Virulence Factor of Pseudomonas aeruginosa. Toxins, 2016. 8(8): p. 236.

59. Diep, B.A., et al., Complete genome sequence of USA300, an epidemic clone of community-acquired meticillin-resistant Staphylococcus aureus. The Lancet, 2006. 367(9512): p. 731-739.

60. Liberati, N.T., et al., An ordered, nonredundant library of Pseudomonas aeruginosa strain PA14 transposon insertion mutants. Proc Natl Acad Sci U S A, 2006. 103(8): p. 2833–8.

61. Holloway, B.W., V. Krishnapillai, and A.F. Morgan, Chromosomal genetics of Pseudomonas. Microbiol Rev, 1979. 43(1): p. 73–102.

62. Bradley, D.E., A study of pili on Pseudomonas aeruginosa. Genet Res (Camb), 1972. 19(1): p. 39–51.

63. Essar, D.W., et al., Identification and characterization of genes for a second anthranilate synthase in Pseudomonas aeruginosa: interchangeability of the two anthranilate synthases and evolutionary implications. J Bacteriol, 1990. 172(2): p. 884–900.

64. Egorova, D.A., et al., Biofilm matrix proteome of clinical strain of P. aeruginosa isolated from bronchoalveolar lavage of patient in intensive care unit. Microb Pathog, 2022. 170: p. 105714.

65. Elmassry, M.M., et al., *Malonate utilization by Pseudomonas aeruginosa affects quorum*-*sensing and virulence and leads to formation of mineralized biofilm*-*like structures*. Molecular microbiology, 2021. 116(2): p. 516–537.

66. Elmassry, M.M., et al., Pseudomonas aeruginosa alters its transcriptome related to carbon metabolism and virulence as a possible survival strategy in blood from trauma patients. Msystems, 2019. 4(4): p. 10.1128/msystems.00312-18.

67. Sauer, K., et al., Characterization of Nutrient-Induced Dispersion in Pseudomonas aeruginosa PAO1 Biofilm. Journal of Bacteriology, 2004. 186(21): p. 7312.

68. She, P., et al., Effects of exogenous glucose on Pseudomonas aeruginosa biofilm formation and antibiotic resistance. MicrobiologyOpen, 2019. 8(12): p. e933.

69. Pisithkul, T., et al., Metabolic Remodeling during Biofilm Development of Bacillus subtilis. mBio, 2019. 10(3): p. e00623–19.

70. Raneri, M., et al., Pseudomonas aeruginosa mutants defective in glucose uptake have pleiotropic phenotype and altered virulence in non-mammal infection models. Scientific Reports, 2018. 8(1): p. 16912.

71. Wang, C., et al., Carbon Starvation Induces the Expression of PprB-Regulated Genes in Pseudomonas aeruginosa. Applied and Environmental Microbiology, 2019. 85(22): p. e01705–19.

72. Dolan, S.K., et al., Contextual Flexibility in Pseudomonas aeruginosa Central Carbon Metabolism during Growth in Single Carbon Sources. mBio, 2020. 11(2): p. e02684–19.

73. Frimmersdorf, E., et al., How Pseudomonas aeruginosa adapts to various environments: a metabolomic approach. Environmental Microbiology, 2010. 12(6): p. 1734–1747.

74. Sauvage, S., et al., Impact of Carbon Source Supplementations on. J Proteome Res, 2022. 21(6): p. 1392–1407.

75. Perinbam, K., et al., A Shift in Central Metabolism Accompanies Virulence Activation in Pseudomonas aeruginosa. mBio, 2020. 11(2).

76. Cornelis, P., Unexpected interaction of a siderophore with aluminum and its receptor. J Bacteriol, 2008. 190(20): p. 6541–3.

77. Camus, L., et al., Trophic cooperation promotes bacterial survival of Staphylococcus aureus and Pseudomonas aeruginosa. The ISME Journal, 2020. 14(12): p. 3093–3105.

78. Hall, S., et al., Cellular effects of pyocyanin, a secreted virulence factor of Pseudomonas aeruginosa. Toxins, 2016. 8: p. 1–14.

79. Miller, J.H., Experiments in molecular genetics. 1972, Cold Spring Harbor, NY: Cold Spring Harbor Laboratory. 466.

80. Schaber, J.A., et al., Analysis of quorum sensing-deficient clinical isolates of Pseudomonas aeruginosa. J Med Microbiol, 2004. 53(Pt 9): p. 841–853.

81. Kharadi, R.R. and G.W. Sundin, Physiological and microscopic characterization of cyclic-di-GMP-mediated autoaggregation in Erwinia amylovora. Front Microbiol, 2019. 10: p. 468.

82. Mahmud, H.A., et al., Synthesis and activity of BNF15 against drug-resistant Mycobacterium tuberculosis. Future Medicinal Chemistry, 2021. 13(03): p. 251–267.

83. Rahme, L.G., et al., Common virulence factors for bacterial pathogenicity in plants and animals. Science, 1995. 268(5219): p. 1899-1902.

84. Takeya, K. and K. Amako, A rod-shaped Pseudomonas phage. Virology, 1966. 28(1): p. 163–165.

85. Fey, P.D., et al., A Genetic Resource for Rapid and Comprehensive Phenotype Screening of Nonessential Staphylococcus aureus Genes. mBio, 2013. 4(1): p. e00537–12.

