## Supplemental Materials for "Global stress response in *Pseudomonas aeruginosa* upon malonate utilization"

**Supplementary Information**

**Supplementary Figures**

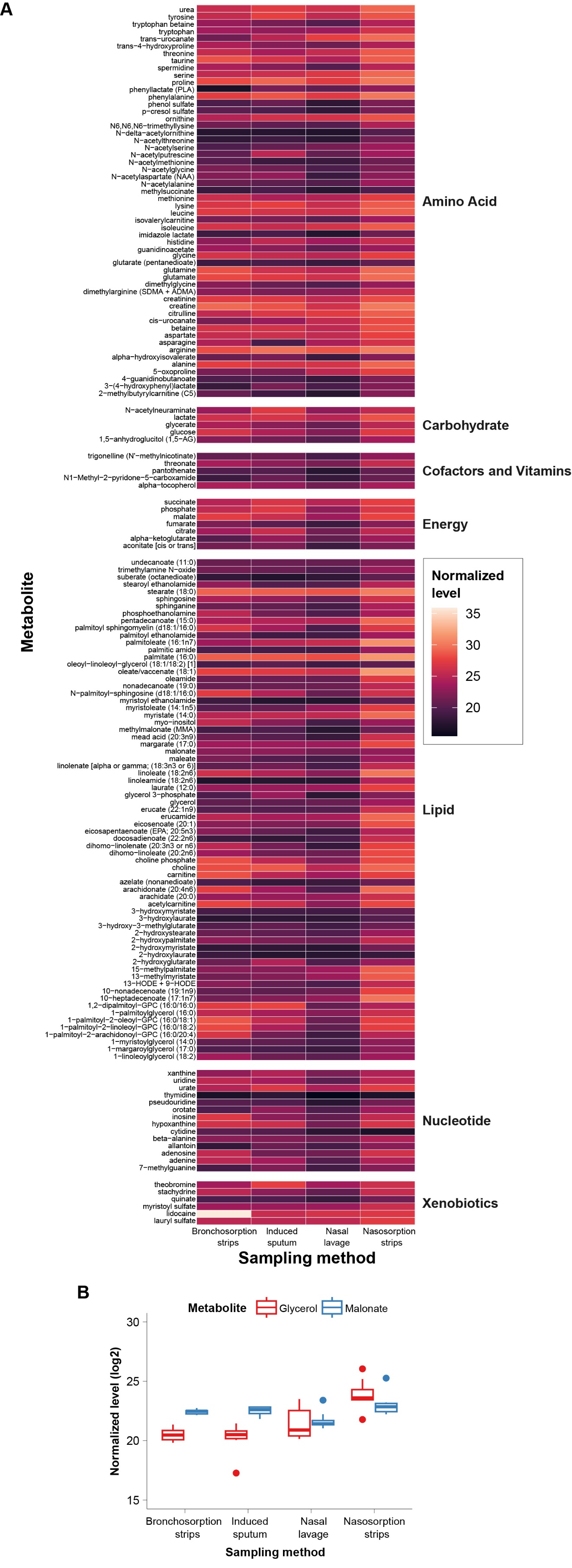
**Supplementary Figure 1. Metabolomic analysis of human airways. (A)** Abundance of 160 metabolites detected in all 32 airway samples and confirmed using chemical standards. **(B)** Abundance of glycerol and malonate across all samples using the four sampling methods used by Farne et al. Data of panel B were extracted from panel A for legibility. Data for both panels were extracted from a publicly available dataset (Farne et al. 2017).

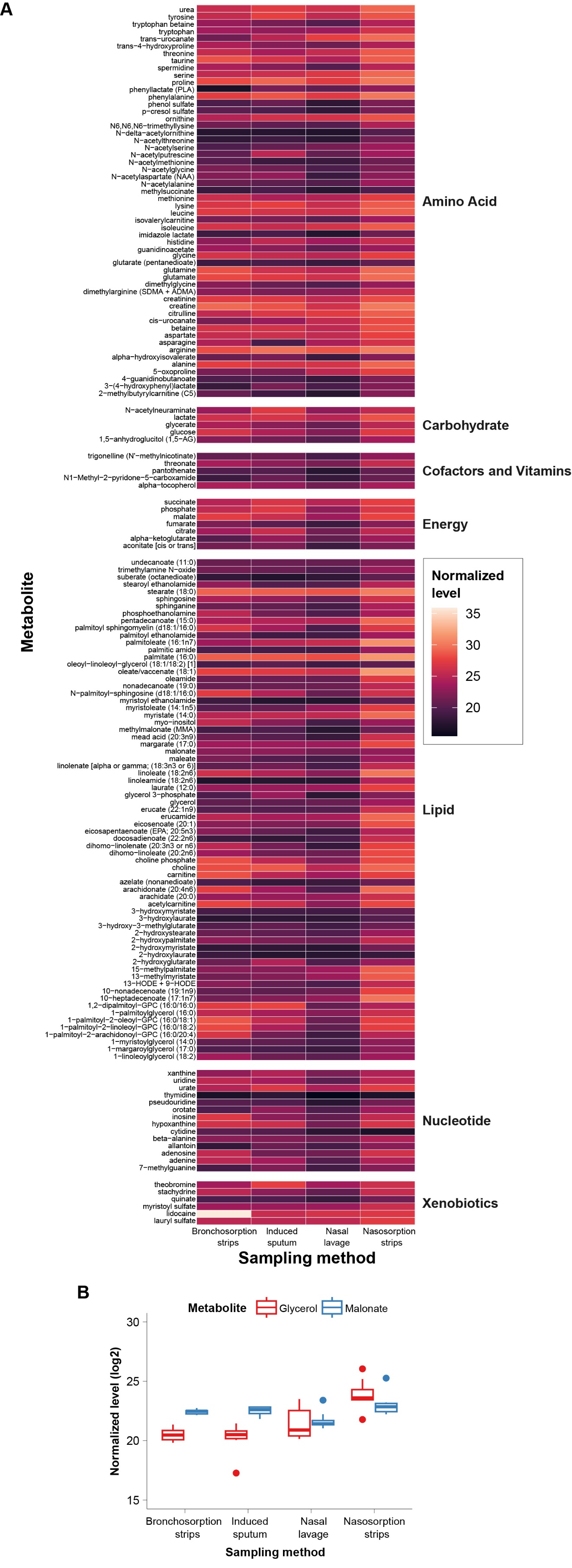

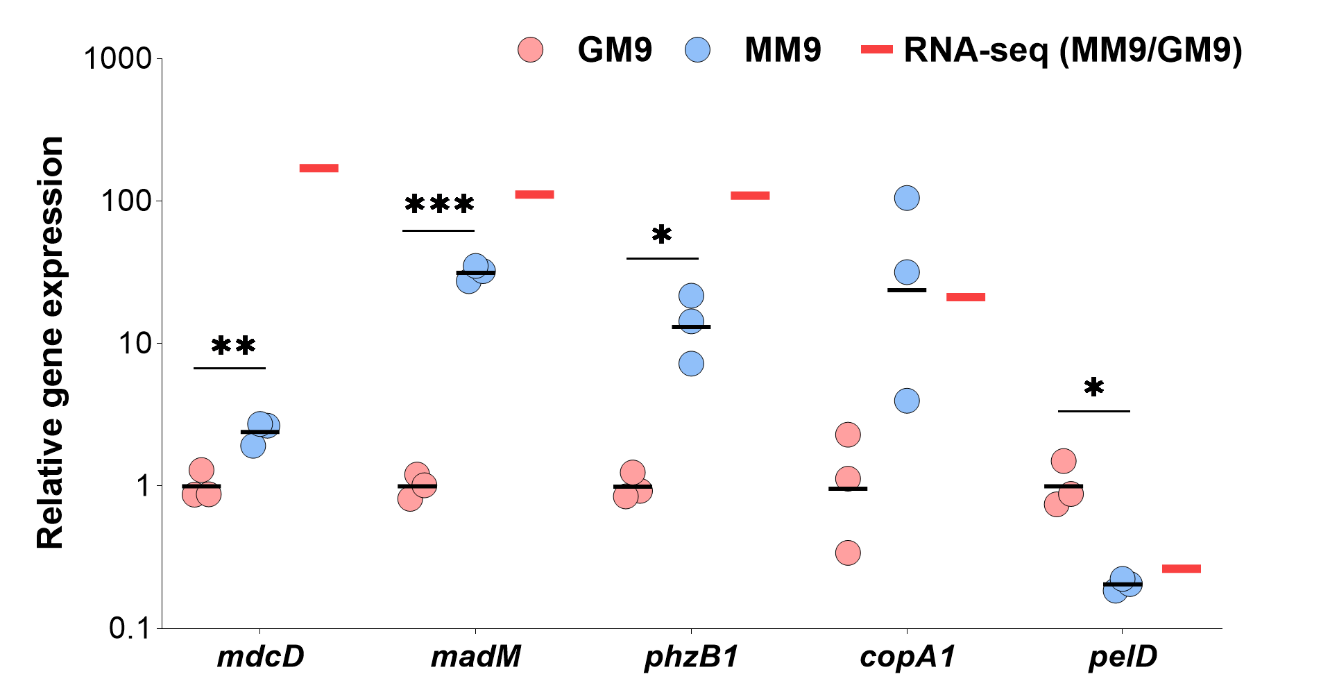

**Supplementary Figure 2. qRT-PCR analysis validates the RNA-seq results.** The 16S ribosomal RNA gene *PA14_08570* was used as a reference gene (housekeeping gene). Horizontal black bars the average of the biological replicates. Red bars represent the expression ratio from the RNA-seq results of MM9 to GM9. Unpaired t-test (two-tailed) was used to measure statistical significance. *P < 0.05, **P < 0.01, and ***P < 0.001.

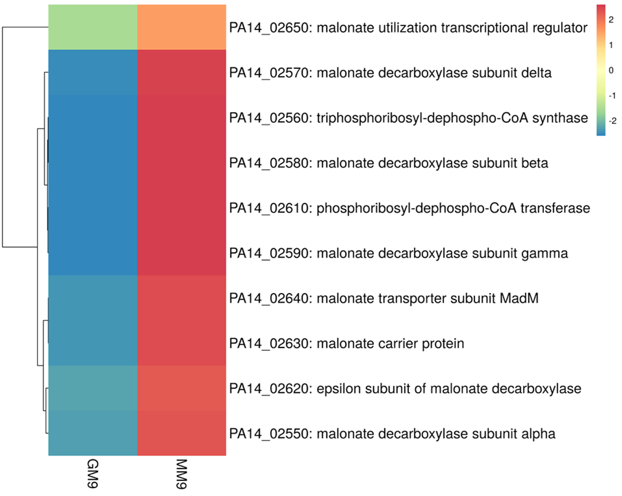

**Supplementary Figure 3. Heatmap of genes involved in malonate uptake and decarboxylation.** All genes are upregulated when *P. aeruginosa* is grown in a minimal medium with malonate as a sole carbon source (MM9) vs. glycerol (GM9). Expression values were pre-processed before plotting by natural log transformation and row centering. Rows are clustered using Euclidean distance and average linkage. Data are presented as average of biological triplicates. Gradient scale is representing expression levels with red showing highest expression and blue showing lowest expression.

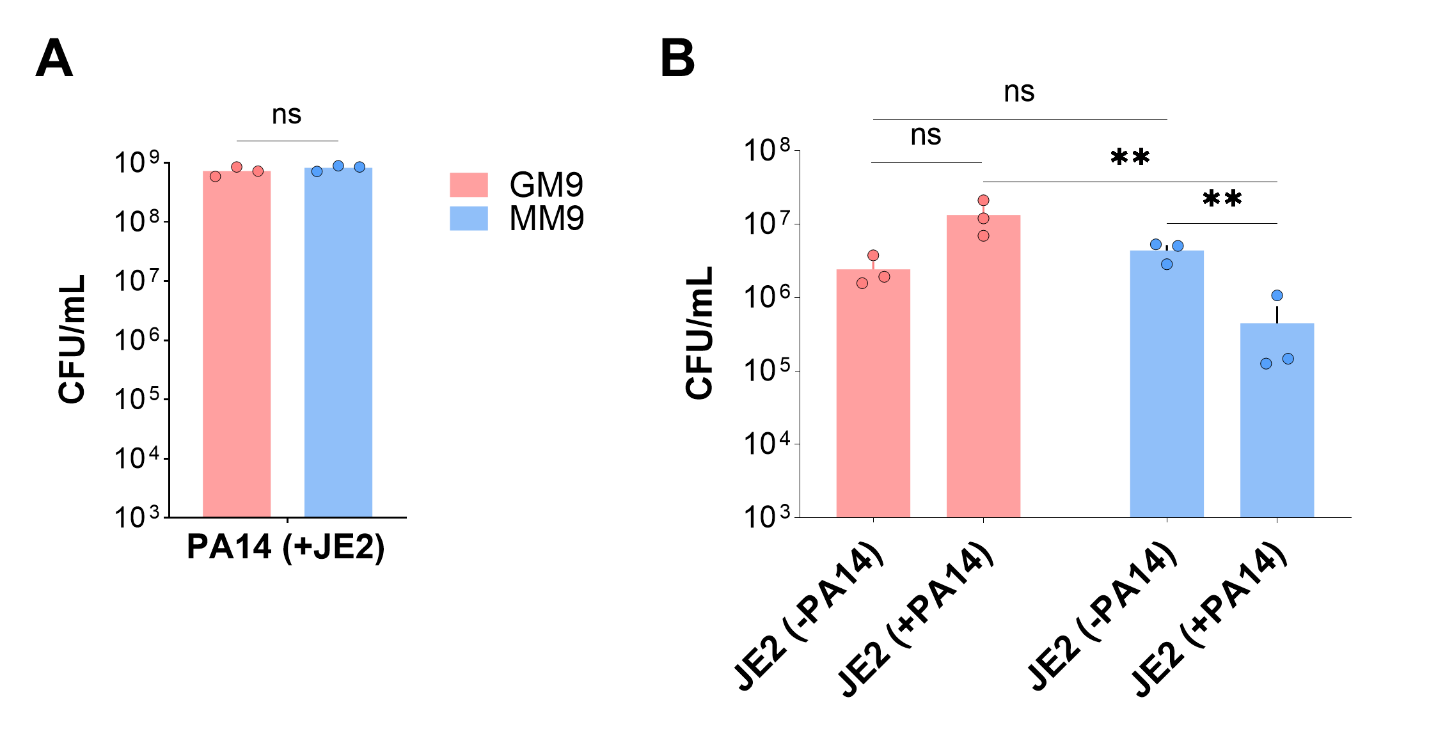

**Supplementary Figure 4. Malonate enhances *S. aureus* JE2 killing by *P. aeruginosa* PA14 in a co-culture condition.** Bacterial abundance of PA14 and JE2 in co-culture in MM9 and GM9 were calculated by counting the colony forming unit (CFU) after cells were serially diluted and plated on selective media, *Pseudomonas* Isolation Agar (PIA) for *P. aeruginosa* and Mannitol Salt Agar (MSA) for *S.* *aureus.* **(A)** No difference was observed in the bacterial abundance of PA14 in co-culture with JE2 in both MM9 and GM9. **(B)** Relative bacterial abundance of JE2 in co-culture with PA14 was significantly low in the presence of MM9 when compared to its abundance in GM9 media. Error bar displays the average from three biological, each with three technical replicates. Error bars indicate SD. Statistical significance was calculated by two-tailed unpaired t-test. ns: not significant, ** P < 0.01.

**Supplementary Tables**

**Table S1.** Selected mutants for growth requirement analysis in MM9 and GM9. PA14 with *MAR2xT7 mariner* transposon insertion within the specific gene. All mutant strains are gentamicin-resistant.

| Gene locus | Gene Name | Gene Description |
| --- | --- | --- |
| PA14_61000 | | conserved hypothetical protein |
| PA14_06750 | *nirS* | nitrite reductase precursor |
| PA14_29850 | *nuoN* | NADH dehydrogenase I chain N |
| PA14_09470 | *phzB1* | probable phenazine biosynthesis protein |
| PA14_13170 | | probable metal transporting P-type ATPase |
| PA14_18070 | | putative periplasmic metal-binding protein |
| PA14_60700 | *ccpR* | cytochrome c551 peroxidase |
| PA14_35160 | | hypothetical protein |
| PA14_01300 | *coxA* | cytochrome c oxidase, subunit I |
| PA14_06870 | *dnr* | transcriptional regulator Dnr |
| PA14_42860 | | putative hemerythrin |
| PA14_66460 | | putative universal stress protein |
| PA14_10550 | | putative sulfite or nitrite reductase |
| PA14_18850 | | putative lyase |
| PA14_44350 | | putative cytochrome c oxidase subunit |
| PA14_18820 | | hypothetical protein |
| PA14_68301 | *arcD* | arginine/ornithine antiporter |
| PA14_11270 | *oprG* | outer membrane protein OprG precursor |
| PA14_36330 | *hcnA* | hydrogen cyanide synthase HcnA |
| PA14_09400 | *phzS* | flavin-containing monooxygenase |
| PA14_29860 | *nuoM* | NADH dehydrogenase I chain M |
| PA14_53070 | *hpd* | 4-hydroxyphenylpyruvate dioxygenase |
| PA14_22320 | | putative membrane protein |
| PA14_53300 | | probable alkyl hydroperoxide reductase |
| PA14_53470 | | probable acetate kinase |
| PA14_09150 | *katA* | catalase |
| PA14_27480 | *htpX* | heat shock protein HtpX |

**Table S2.** List of primers used in this study.

| Gene and primer direction | Sequence |
| --- | --- |
| mdcD_F | 5'-GTACGCACCTCGGTACTCGC-3' |
| mdcD_R | 5'-ATGTTGCATCTCCTTGGCCC-3' |
| madM_F | 5'-CATGGCGGTGTCCTACTGGT-3' |
| madM_R | 5'-CACCAATCCTTTCTGCCCGC-3' |
| phzB1_F | 5'-TAAACCGCCACAGCCATCCT-3' |
| phzB1_R | 5'-GGTTTCACCGATGCCAACGAA-3' |
| copA1_F | 5'-GCCAGAGGTAGACGCTGAGG-3' |
| copA1_R | 5'-GCGCGCGCTTCTATGTCTC-3' |
| pelD_F | 5'-TTTCCCTGGGTGATCCTCGC-3' |
| pelD_R | 5'GAACTCGCCCACCAACATGG-3' |
| PA14_21220_F | 5'-CCTGGAAGAGGCCATCAAG-3' |
| PA14_21220_R | 5'-GCTCGCCTCGACCTTCTC-3' |
| PA14_18070_F | 5'-TGCAAGTCTTCAAGGTTCAGG-3' |
| PA14_18070_R | 5'-CGGATCGCCTCGAGTACCT-3' |
| PA14_09150_F | 5'-CGGCAAGAAGACCGATATGT-3' |
| PA14_09150_R | 5'-GTTGAGATCGGGGAACTTGA-3' |
| PA14_68300_F | 5'-TTCGTCCTGTTGTTCAGCAC-3' |
| PA14_68300_R | 5'-GGCAGATCAGGATGAACAGG-3' |
| PA14_27480_F | 5'-AGCATCACCCTGAAACTGCT-3' |
| PA14_27480_R | 5'-AGCAGCCACTGTTCGTGAC-3' |
| PA14_62990_F | 5'-GGATGGAACTGACCCTGAAG-3' |
| PA14_62990_R | 5'-CGTTGAGCAGGTAGCCTTTC-3' |
| PA14_08570_F | 5’-AAGCAACGCGAAGAACCTTA-3’ |
| PA14_08570_R | 5’-CACCGGCAGTCTCCTTAGAG-3’ |
